# scRNA-seq and genomics analyses reveal key mechanisms of inverted papilloma-associated sinonasal squamous cell carcinoma malignant transformation

**DOI:** 10.64898/2026.06.01.729345

**Authors:** Tolani F. Olonisakin, Fernando T. Zamuner, Kenny Vu, Sreenivasulu Gunti, Yiming Ding, Yue Hou, Xinping Yang, Ramya Viswanathan, Angel Huynh, Federico Comoglio, Marco Notaro, Christopher Cherry, Michael Patatanian, Nathan Wong, Timothy R. DeKlotz, Jean Kim, Wayne Koch, Andrew P. Lane, Murugappan Ramanathan, Nicholas R. Rowan, Matt Lechner, Aaron Udager, Evgeny Izumchenko, Clint T. Allen, Nyall R. London

## Abstract

Sinonasal inverted papilloma (IP) carries a 10% risk of malignant transformation to IP-associated sinonasal squamous cell carcinoma (IP-SNSCC), yet the molecular and immune drivers of this progression remain poorly defined. This study integrates single-cell RNA sequencing, multiplexed spatial proteomics, whole-exome sequencing, and functional assays across IP and IP-SNSCC cohorts to define mechanisms of malignant transformation. Respiratory epithelial basal cells are identified as the putative cell of origin, with progression marked by recurrent *CDKN2A* loss and *TP53* mutations. Spatial profiling reveals immune reorganization at the lesional interface in IP-SNSCC, characterized by enrichment of alternatively activated M2 macrophages. CXCL14 is shown to directly induce an immunosuppressive myeloid phenotype that suppresses T-cell IFNγ production through an IDO-pathway-dependent mechanism. Integration of these multimodal datasets defines a previously unrecognized CXCL14-IDO mechanism that constrains anti-tumor immunity at the tumor-stroma interface. These findings establish IDO-targeted immunomodulation as a rational adjuvant strategy and provide a comprehensive molecular framework for understanding IP-SNSCC pathogenesis.

## Introduction

Sinonasal squamous cell carcinoma (SNSCC) is the most common sinonasal malignancy, accounting for ∼60-75% of sinonasal cancers^1–3^. Despite improvements in outcomes for patients with squamous cell carcinomas arising at other head and neck cancer subsites over the last three decades, the prognosis for patients with SNSCC has remained largely unchanged, with 5-year overall survival hovering around 50%^1^. This stagnation reflects a limited understanding of the biological mechanisms underlying SNSCC pathogenesis. Surgical resection with curative intent remains the primary treatment modality, and although advances in endoscopic approaches have reduced treatment-related morbidity^4^, surgery is frequently constrained by the close proximity of sinonasal tumors to critical neurovascular structures^1^. Consequently, there is a pressing need for effective adjunct and alternative therapeutic strategies.

Sinonasal inverted papilloma (IP) is one of the most frequently observed benign sinonasal neoplasms, with malignant transformation to IP-derived sinonasal squamous cell carcinoma (IP-SNSCC) occurring in up to 10% of patients^5,6^. The mechanisms underlying progression from IP to IP-SNSCC remain poorly understood, and molecular features that distinguish IP-SNSCC from IP are incompletely defined. *EGFR* exon 20 activating mutations or low-risk human papillomavirus (HPV) serotypes have been detected in both IP and IP-SNSCC tumors^7–9^, supporting the concept that these lesions represent a continuum of neoplastic evolution^9^. Moreover, *CDKN2A* mutations have been observed in IP-SNSCC tumors but are absent in matched benign precursor lesions, suggesting that these alterations may represent early molecular events in malignant progression^10^.

Beyond epithelial cell-intrinsic alterations, malignant progression is also likely to depend on remodeling of the local immune microenvironment^11^. Spatially organized myeloid immunoregulatory states, including those shaped by chemokine signaling and metabolic immune checkpoint pathways, may limit productive anti-tumor immunity despite substantial immune infiltration^12,13^. However, the molecular mechanisms governing these local immune states during progression from IP to IP-SNSCC remain unresolved.

Here, we performed the first single-cell RNA sequencing (scRNA-seq) of IP and IP-SNSCC tumors using 10x Genomics 5′ gene expression profiling. We further integrated whole-exome sequencing, spatial proteomic profiling, and functional assays to characterize IP and IP-SNSCC and define molecular and immune features associated with disease progression. Through this multi-modal approach, we identify candidate mechanisms linking malignant transformation to local immunoregulation and, to our knowledge, provide the first integrated molecular analysis of neoplastic transformation from IP to IP-SNSCC.

## Results

### Cohort characteristics

After quality assessment, 21 specimens collected from 19 patients (15 with benign sinonasal papilloma and 4 with IP-SNSCC) were retained for downstream scRNA-seq analyses. WES was performed in 18 patients (14 benign sinonasal papillomas and 4 IP-SNSCC), yielding 36 DNA samples (18 tumor-normal pairs). Seventeen patients were included in both datasets, whereas two patients were represented only in the scRNA-seq cohort and one IP-SNSCC patient only in the WES dataset. Clinicopathologic and demographic characteristics are summarized in **Table 1**. The mean age at diagnosis was 63.9 years (s.d. 10.6), with a median of 69.0 years (interquartile range, 54.0 - 72.0). Patients with benign sinonasal papillomas had a mean age of 63.1 years, compared with 66.2 years for patients with IP-SNSCC. The cohort was predominantly male (15 of 20, 75%), including 11 cases with IP (73.3%) and 4 with IP-SNSCC (80.0%). Self-reported race was White in 60% of patients (12 of 20), Black in 35% (7 of 20), and Asian in 5% (1 of 20). Most patients were never-smokers (13 of 20, 65%), whereas 3 of 20 (15%) were former smokers and 4 of 20 (20%) were current smokers.

**Table 1.**
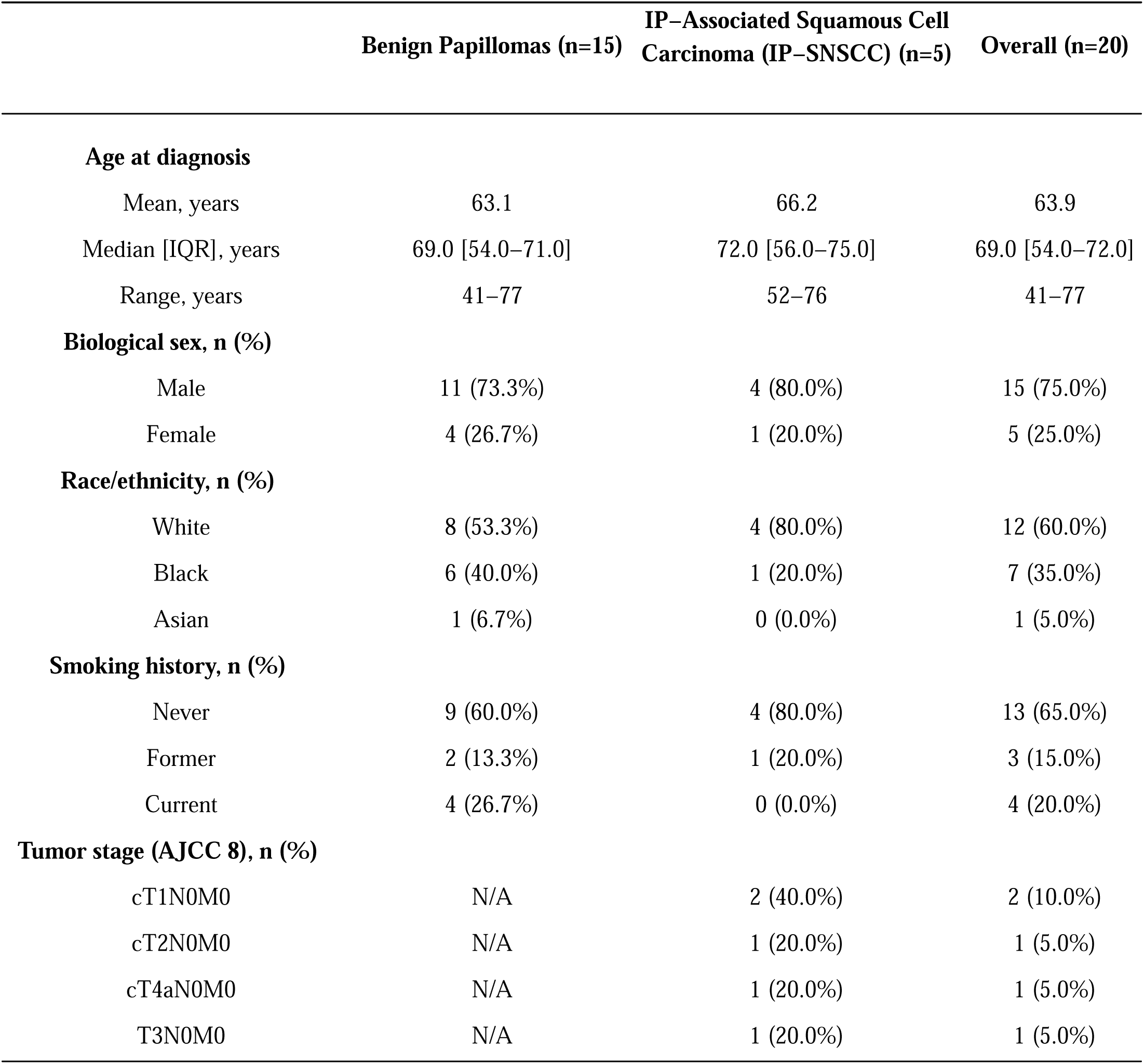
Patient demographics.

|  | Benign Papillomas (n=15) | IP-Associated Squamous Cell Carcinoma (IP-SNSCC) (n=5) | Overall (n=20) |
| --- | --- | --- | --- |
| **Age at diagnosis** |  |  |  |
| Mean, years | 63.1 | 66.2 | 63.9 |
| Median [IQR], years | 69.0 [54.0–71.0] | 72.0 [56.0–75.0] | 69.0 [54.0–72.0] |
| Range, years | 41–77 | 52–76 | 41–77 |
| **Biological sex, n (%)** |  |  |  |
| Male | 11 (73.3%) | 4 (80.0%) | 15 (75.0%) |
| Female | 4 (26.7%) | 1 (20.0%) | 5 (25.0%) |
| **Race/ethnicity, n (%)** |  |  |  |
| White | 8 (53.3%) | 4 (80.0%) | 12 (60.0%) |
| Black | 6 (40.0%) | 1 (20.0%) | 7 (35.0%) |
| Asian | 1 (6.7%) | 0 (0.0%) | 1 (5.0%) |
| **Smoking history, n (%)** |  |  |  |
| Never | 9 (60.0%) | 4 (80.0%) | 13 (65.0%) |
| Former | 2 (13.3%) | 1 (20.0%) | 3 (15.0%) |
| Current | 4 (26.7%) | 0 (0.0%) | 4 (20.0%) |
| **Tumor stage (AJCC 8), n (%)** |  |  |  |
| cT1N0M0 | N/A | 2 (40.0%) | 2 (10.0%) |
| cT2N0M0 | N/A | 1 (20.0%) | 1 (5.0%) |
| cT4aN0M0 | N/A | 1 (20.0%) | 1 (5.0%) |
| T3N0M0 | N/A | 1 (20.0%) | 1 (5.0%) |

### Distinct cellular landscape of the IP and IP-SNSCC tumor microenvironment

To uncover the molecular mechanisms driving malignant transformation, we performed the first known scRNA-seq analysis of tumors from patients with IP and IP-SNSCC (**Fig. 1a**). Unbiased clustering was applied to the combined single cell transcriptomes, identifying 25 distinct cell clusters (**Fig. 1b**). Although cluster proportions varied, each cluster was consistently observed across tumor samples rather than reflecting patient-specific subpopulations (**Fig. S1a-d**). Differential expression analysis revealed canonical marker genes specific to each cluster, delineating major cell populations and subpopulations, including ciliated epithelial cells, malignant epithelial cells, endothelial cells, fibroblasts, pericytes, T cells, B cells, mast cells, and myeloid cells (**Fig. 1c**). We next performed spatial phenotyping on formalin-fixed paraffin-embedded (FFPE) tissue from IP and IP-SNSCC tumors using the PhenoCycler-Fusion system (**Fig. 1d**). A panel of 25 markers was applied (**Fig. S2a**), and single-cell segmentation was carried out using StarDist to define nuclear and cell membrane boundaries (**Fig. S2b-d**). IP-SNSCC tumors were relatively immune-rich compared to IP neoplasms (**Fig. 1e, f**), displaying a diverse immune infiltrate (**Fig. 1g**), consistent with prior reports describing sinonasal squamous cell carcinoma as an immunogenic tumor type^14,15^.

**Figure 1.**
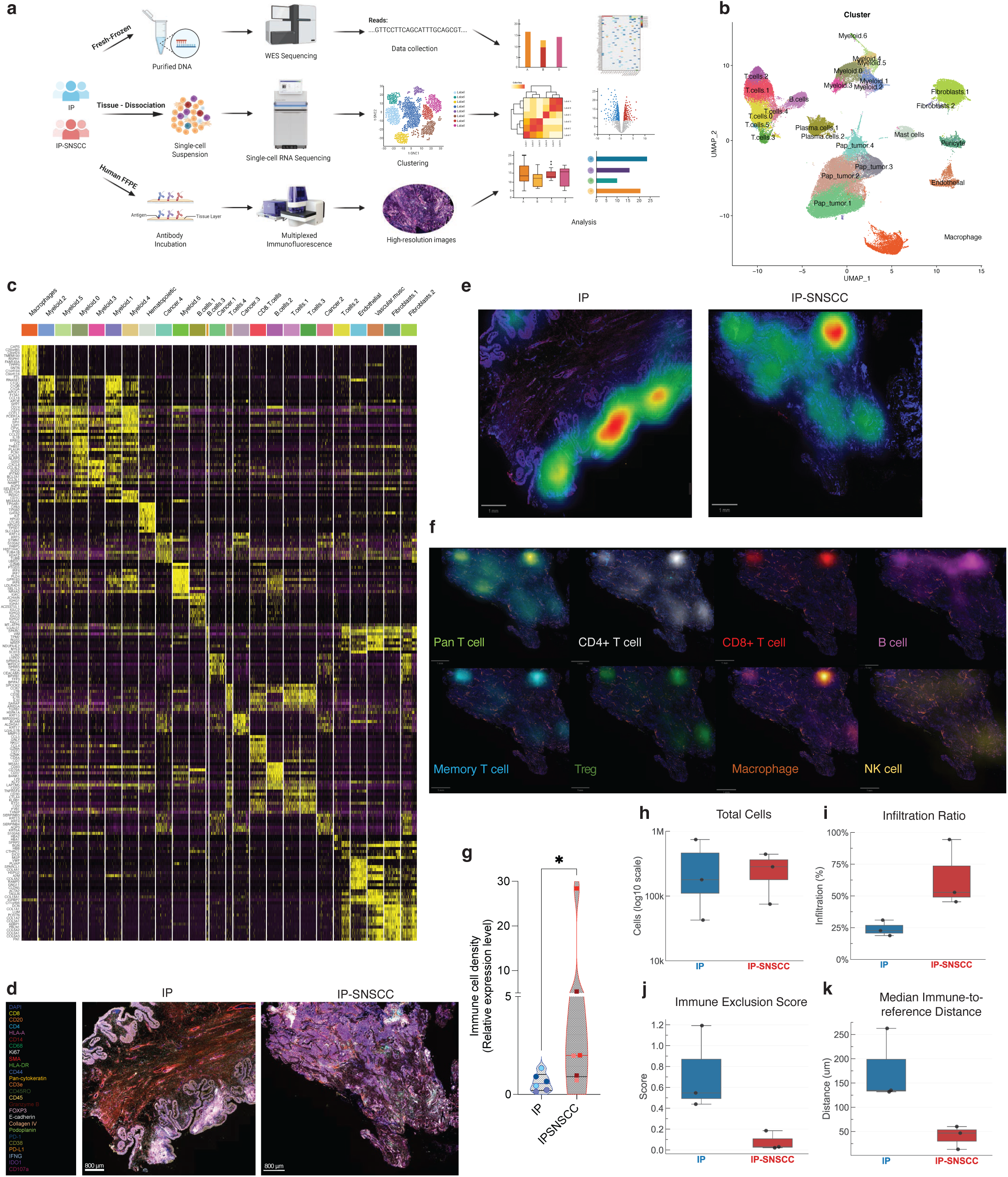
Distinct cellular and spatial organization of the IP and IP-SNSCC tumor microenvironment. a, Study schematic showing the integrated analytical workflow. Fresh-frozen IP and IP-SNSCC specimens were dissociated for single-cell RNA sequencing and whole-exome sequencing, and FFPE tissues were analyzed by multiplexed immunofluorescence followed by image-based spatial analysis. **b,** Uniform manifold approximation and projection (UMAP) of single-cell transcriptomes showing the major epithelial, immune and stromal populations identified across the cohort. **c,** Heat map of the top differentially expressed genes across annotated cell clusters. **d,** Representative multiplexed immunofluorescence images from IP and IP-SNSCC specimens. **e,** Immune-density hotspot maps in representative IP and IP-SNSCC sections. **f,** Spatial maps showing the distribution of major immune populations, including pan-T cells, CD4^+^ T cells, CD8^+^ T cells, B cells, memory T cells, regulatory T cells, macrophages and natural killer cells. **g,** Violin plot showing CD45^+^ cell density per μm^2^ in IP and IP-SNSCC tumors. n = 3 biological replicates per group, with 2 tissue sections analyzed per patient; two-sided Mann-Whitney test, P < 0.05. Solid lines indicate the median. **h,** Total number of segmented cells identified per specimen from PhenoCycler-Fusion images. **i,** Immune infiltration ratio, defined as the fraction of immune cells located within 50 μm of the lesional compartment. **j,** Immune exclusion score, calculated as the fraction of immune cells located >200 μm from the lesional compartment divided by the fraction located within 0-200 μm. **k,** Median distance from immune cells to the lesional compartment for each specimen.

### Immune cells are spatially reorganized toward the lesional compartment in IP-SNSCC

To define immune cell positioning relative to the lesional compartment, we measured the distance from each immune cell to its nearest reference-compartment cell (tumor in IP-SNSCC and epithelium in IP). The mean infiltration ratio (the fraction of immune cells located within 50 µm of the reference compartment) was higher in IP-SNSCC than in IP (64.2% versus 24.1%), whereas the mean immune exclusion score was strikingly lower (0.08 versus 0.73) and the median immune-to-reference distance was markedly shorter (40.1 µm versus 176.5 µm) (**Fig. 1g, i-k**). The core-to-far ratio, defined as the number of immune cells within 50 µm of the lesional compartment relative to those located beyond 200 µm, was 22.5 in IP-SNSCC compared with 0.61 in IP, corresponding to an approximately 37-fold difference. Notably, all three IP-SNSCC specimens exhibited lower median immune-to-reference distances than any IP specimen, yielding complete rank separation between cohorts for this metric (**Fig. 1k**). By contrast, total cell burden overlapped across groups (IP-SNSCC, 75,090-442,609 cells; IP, 42,618-745,225 cells), indicating that the defining distinction between cohorts lies in immune cell spatial organization relative to the lesional compartment rather than in overall specimen cellularity (**Fig. 1h**).

### Pseudotime trajectory analysis uncovers basal cells of the respiratory epithelium as the putative cell of origin for IP development

Next, we integrated publicly available scRNA-seq data obtained from healthy sinonasal mucosa brushings and biopsies^16^ to generate a cluster map combining normal, benign sinonasal papilloma and IP-SNSCC transcriptomes. Consistent with the analysis restricted to IP and IP-SNSCC samples, the integrated dataset again resolved into 25 analogous cell clusters (**Fig. 2a**). When examined separately, normal sinonasal mucosa was characterized by diverse epithelial subtypes, including multiciliated epithelial cells, goblet cells, secretory cells, serous cells and basal cells (**Fig. 2b-c**). Notably, basal cells, rather than ciliated, goblet, secretory or serous cells, mapped to the papilloma cluster within the IP dataset (**Fig. 2c-e**), suggesting that basal cells serve as precursors to squamous cell papilloma. To validate epithelial differentiation trajectories within our dataset, we applied Slingshot analysis to model changes in gene expression and cell ordering along pseudotime. Differential gene expression comparing normal tissue to IP, normal tissue to IP-SNSCC, and IP to IP-SNSCC within each epithelial cluster was largely consistent across clusters (**Fig. S3a-d**). Ciliated epithelial cells appeared earliest along the developmental trajectory and showed the potential to diverge into two distinct lineages (**Fig. 2f**). As cells transitioned from normal respiratory epithelium to papillomatous tumors, we observed progressive downregulation of genes essential for ciliary motility, including *DNAH5*, *DNAH9*, *DNAH12* in both lineage 1 (**Fig. 2g**) and lineage 2 (**Fig. 2h**), accompanied by upregulation of genes associated with stemness and basal epithelial identity such as *KRT14* and *LGALS7*. Furthermore, we observed transient upregulation along the pseudotime trajectory of the type II keratin *KRT80*, inflammatory-associated genes including *BPIFB1* (LPLUNC1) and *C15ORF48* (MOCCI), as well as *TFF3* (a marker of respiratory mucosa goblet cell)^17^, in both lineages (**Fig. 2g, h**). Interestingly, lineage 2 was primarily characterized by upregulation of genes regulating mitosis, namely *UBE2C*, *PLK1*, *TPX2*, *CENPF*, and *ASPM* (**Fig. 2h**), whereas lineage 1 was distinguished by genes involved in extracellular matrix remodeling, including *SERPINE1*, *SERPINE2*, *CEMIP*, *COL17A1*, *MCAM*, and the chemokine *CXCL14* (**Fig. 2g**).

**Figure 2.**
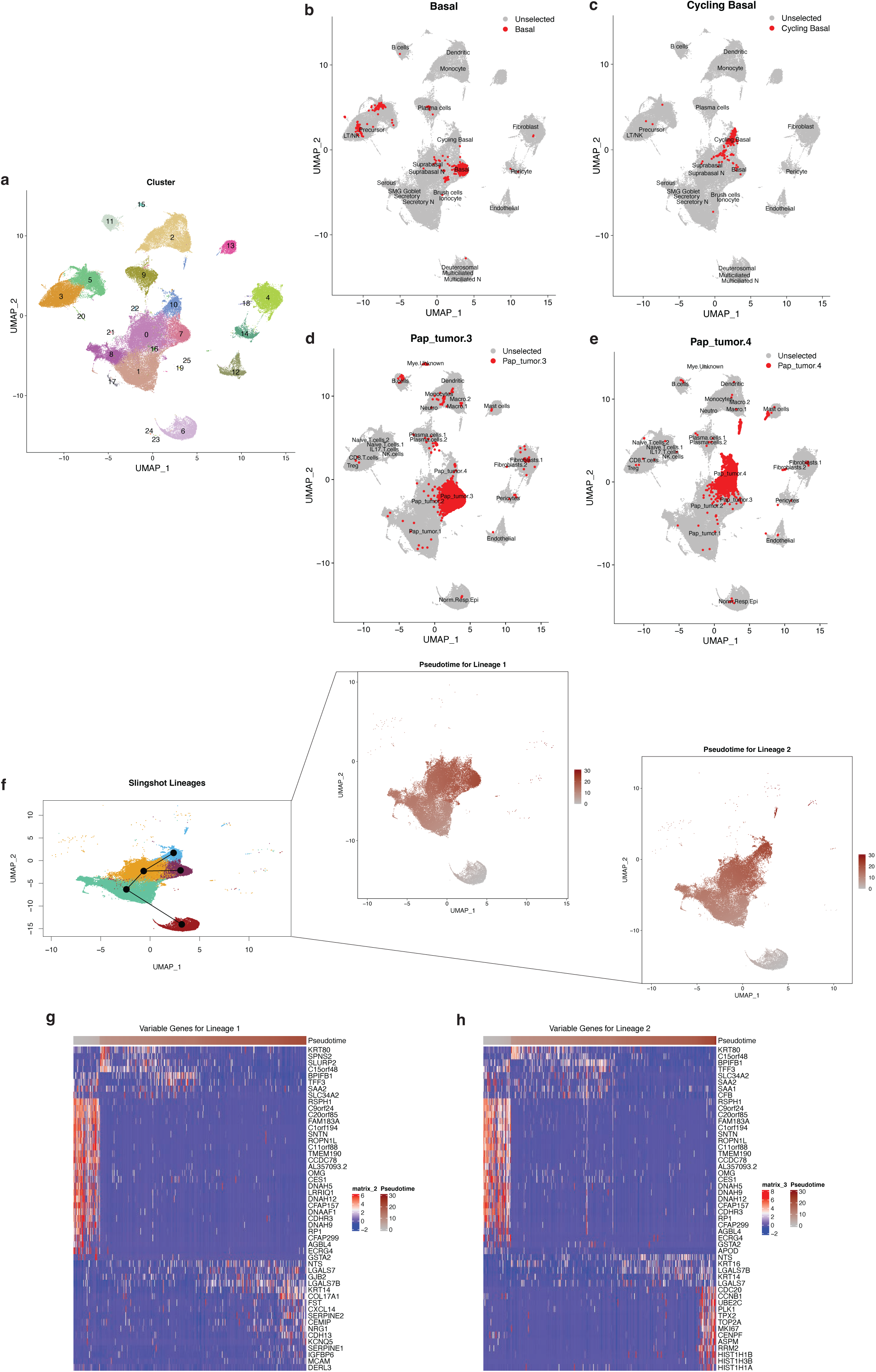
Basal-like epithelial states define putative trajectories of papilloma development and progression. a, Uniform manifold approximation and projection (UMAP) of the integrated epithelial/tumor-cell dataset combining healthy sinonasal mucosa, IP and IP-SNSCC transcriptomes, showing unsupervised clustering of epithelial states. **b,c,** UMAPs highlighting the distribution of basal **(b)** and cycling basal **(c)** epithelial populations within the integrated dataset. **d,e,** UMAPs highlighting the distribution of the papillomatous/malignant epithelial clusters Pap_tumor.3 **(d)** and Pap_tumor.4 **(e).** In **b-e,** selected cells are shown in red and all remaining cells are shown in gray. **f,** Slingshot trajectory inference overlaid on the UMAP, identifying two principal lineages emerging from basal-like epithelial states and extending toward distinct papillomatous/tumor endpoints. Corresponding pseudotime values for lineage 1 and lineage 2 are shown in the adjacent UMAP projections. **g,h,** Heat maps of genes varying along lineage 1 **(g)** and lineage 2 **(h)** as a function of pseudotime. Cells are ordered by inferred pseudotime, and rows represent dynamically expressed genes across each lineage. Color intensity indicates scaled expression.

### Myeloid cells are enriched in the IP-SNSCC tumor microenvironment compared to IP

Given the heterogeneous immune landscape revealed by scRNA-seq profiling, we next interrogated the immune composition of IP and IP-SNSCC neoplasms. Cluster proportion analysis comparing immune cell populations across samples showed a consistent trend towards increased myeloid cell representation in IP-SNSCC tumors (**Fig. 3a-g**), with marked enrichment of myeloid cell cluster 1 relative to sinonasal papillomas (**Fig. 3b**). By contrast, no notable trends or significant differences were observed in the proportions of B cells, mast cells, natural killer cells, CD8+ T cells, or regulatory T cells between IP and IP-SNSCC tumors (**Fig. 3h-l**).

**Figure 3.**
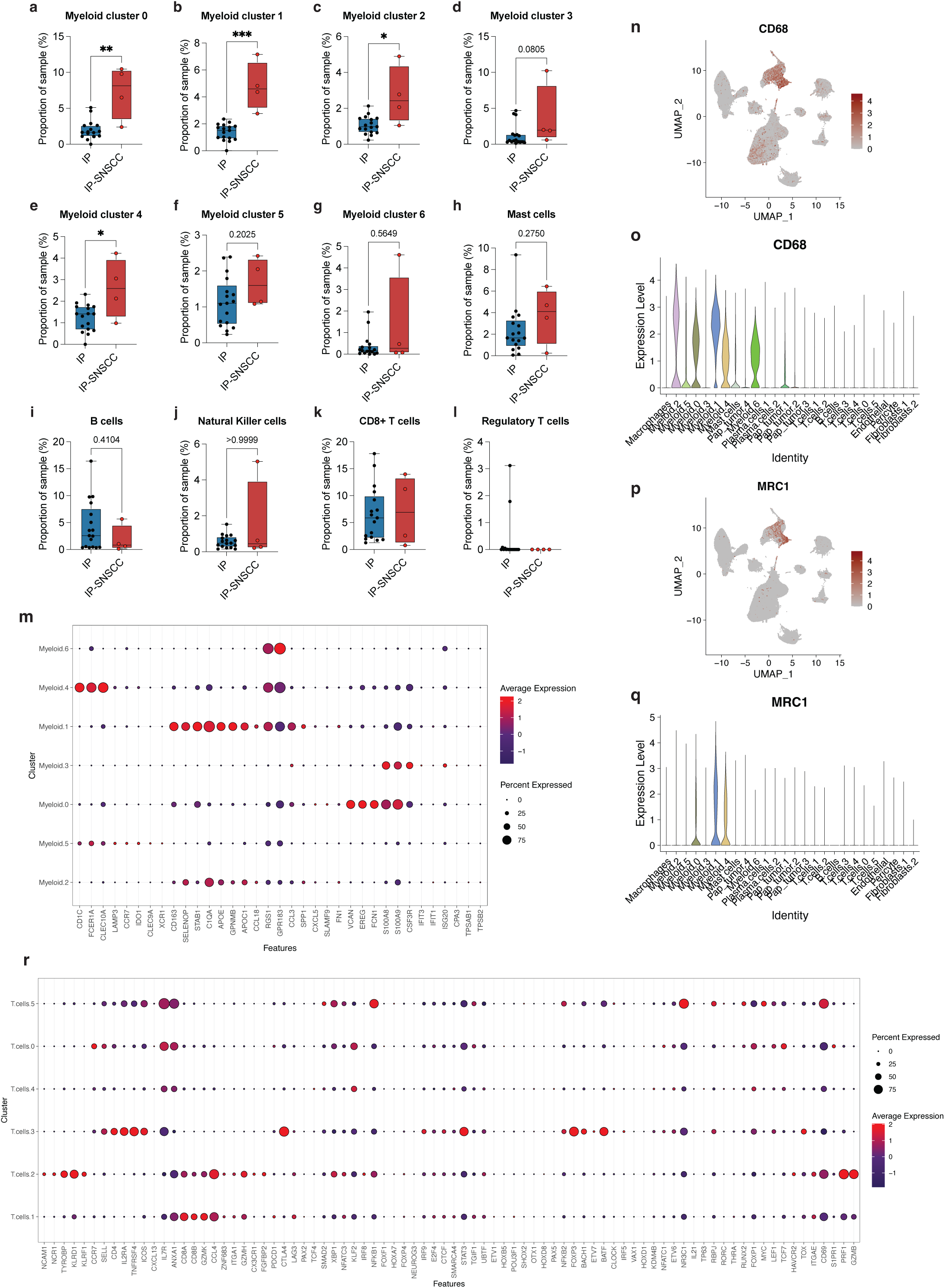
**Single-cell immune profiling identifies myeloid remodeling and a macrophage program marked by CD68 and MRC1 expression in IP-SNSCC. a-g**, Per-sample proportions of myeloid clusters 0-6 in IP and IP-SNSCC. **h-l**, Per-sample proportions of mast cells, B cells, natural killer cells, CD8^+^ T cells and regulatory T cells. **m,** Dot plot showing marker-gene expression across myeloid clusters. Myeloid cluster 1 is characterized by expression of *CD163*, *SELENOP*, *STAB1*, *C1QA*, *APOE*, *CD68* and *MRC1*, consistent with a macrophage program with alternatively activated or immunoregulatory features. Myeloid cluster 4 expresses *CD1C*, *CLEC10A* and *FCER1A*, consistent with a dendritic cell state, whereas clusters 0 and 3 are enriched for *S100A8* and *S100A9*, with cluster 0 additionally expressing *EREG*. **n,o,** Feature and violin plots showing *CD68* expression across immune cell clusters. **p,q,** Feature and violin plots showing *MRC1* expression across immune cell clusters. **r,** Dot plot showing marker-gene expression across T-cell clusters. Box plots show the median and interquartile range, with whiskers extending to 1.5 × the interquartile range; points represent individual samples. In dot plots, dot size indicates the fraction of cells expressing each gene and color indicates scaled average expression.

### IP-SNSCC is enriched for a macrophage program marked by CD68 and MRC1 expression

We further characterized myeloid subpopulations identified in our unsupervised clustering analysis. Interestingly, expression of the G-protein coupled receptor 183 (*GPR183*) and regulator of G-protein signaling 1 (*RGS1*) emerged as a shared feature across multiple myeloid subpopulations (**Fig. 3m**). Expression of the alarmins *S100A8* and *S100A9*, was a defining feature of myeloid cell clusters 0 and 3 (**Fig. 3m**). In addition, myeloid cell cluster 0 was characterized by expression of the epidermal growth factor receptor ligand epiregulin (*EREG*), a feature consistent with a subset of dendritic cells (DC3 dendritic cells)^18^. Classical dendritic cell markers *CD1C* and *CLEC10A* were defining features of myeloid cluster 4, together with *FCER1A* (**Fig. 3m**). The most enriched myeloid cell cluster (myeloid cluster 1) was characterized by upregulation of *CD163*, *SELENOP*, *STAB1*, *C1QA*, and *APOE*. (**Fig. 3m**), and further defined by expression of *CD68* and *MRC1* (*CD206*), established markers of alternatively activated or immunosuppressive “M2” macrophages^19^ (**Fig. 3n-q**). Notably, *MRC1*/*CD206* expression in this subset did not colocalize with dendritic (*ITGAX*, **Fig. S4a**) or granulocytic (*CSF3R*, **Fig. S4b**) markers, supporting classification of cluster 1 as a macrophage population rather than a dendritic or granulocytic subset. T-cell subclustering identified diverse lymphoid states, with no evidence of a dominant IP-SNSCC-enriched T-cell program (**Fig. 3r**).

### Macrophages are positioned closer to the lesional interface in IP-SNSCC

We next assessed whether this interface-proximal spatial redistribution (defined by increased immune infiltration ratio, reduced immune exclusion score, shorter median immune-to-reference distance, and higher core-to-far infiltration ratio) extended specifically to the macrophage compartment. In an exploratory analysis, macrophages in IP-SNSCC were positioned closer to the reference compartment than those in IP, with a higher macrophage infiltration ratio (cohort median, 65.4% versus 34.6%) and a shorter median macrophage-to-reference distance (32.3 µm versus 103.2 µm) (**Fig. 4a, b**). Notably, all three IP-SNSCC specimens exhibited higher macrophage infiltration than all IP specimens, yielding complete rank separation between the two cohorts. Consistent with this pattern, macrophages in IP-SNSCC were concentrated within the core (0-50 µm) zone, whereas macrophages in IP lesions were distributed more broadly across the margin and far interface-distal zones (**Fig. 4c**). This shift was further reflected in a higher core-to-far infiltration score in IP-SNSCC (**Fig. 4d**). Together, these findings support interface-proximal macrophage organization in IP-SNSCC.

**Figure 4.**
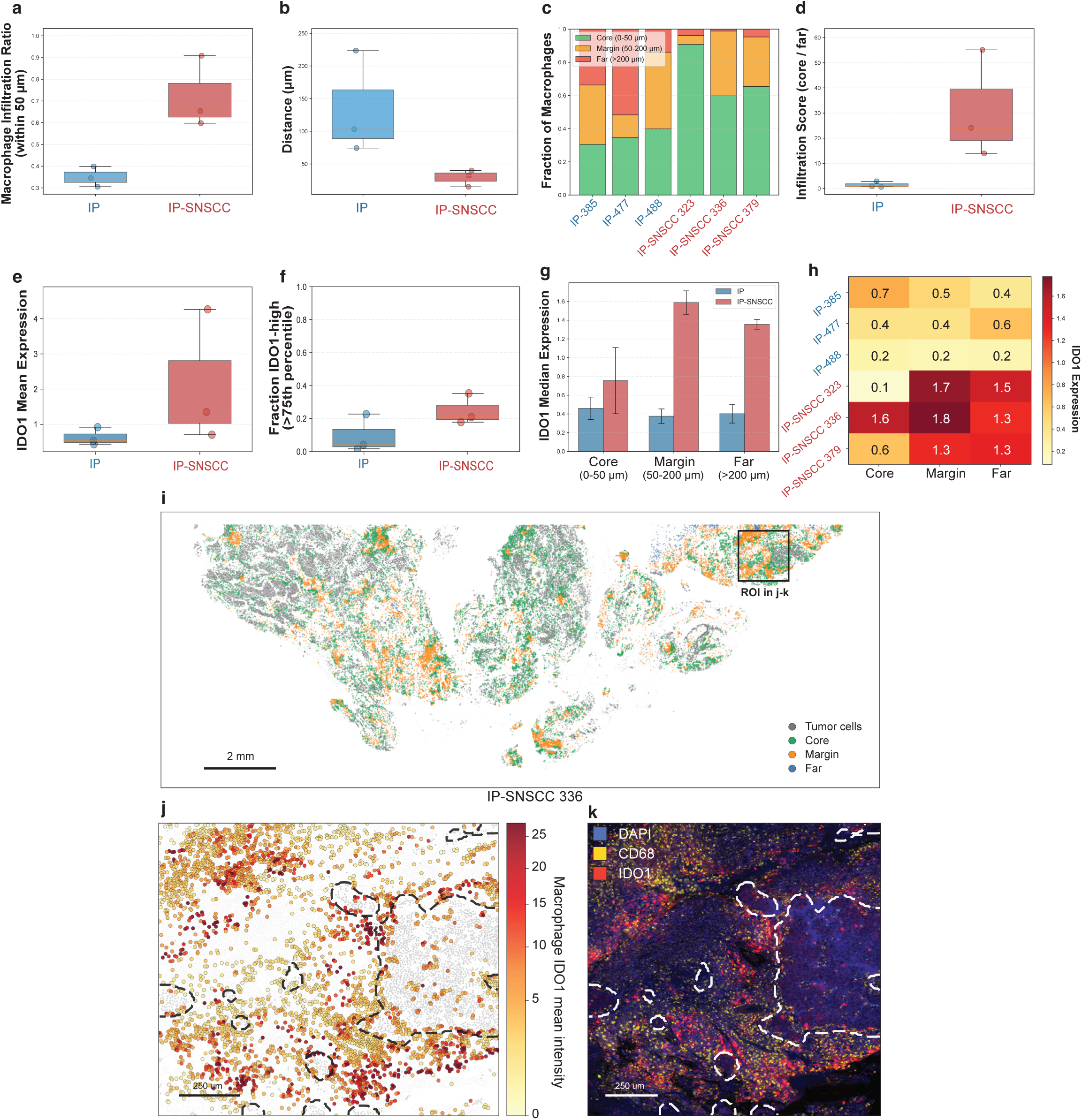
Macrophages in IP-SNSCC localize to the lesional interface and exhibit spatially patterned IDO1 expression. a, Macrophage infiltration ratio, defined as the fraction of macrophages located within 50 µm of the reference compartment. **b,** Median macrophage-to-reference distance. **c,** Fraction of macrophages distributed across core (0-50 µm), margin (50-200 µm) and far (>200 µm) zones relative to the reference compartment for each sample. **d,** Core-to-far infiltration score. **e,** Mean IDO1 expression in macrophages by group. **f**, Fraction of IDO1-high macrophages by group, using a >75th percentile threshold. **g,** Median macrophage IDO1 expression stratified by spatial zone in IP and IP-SNSCC. **h,** Heat map showing macrophage IDO1 expression by sample and spatial zone. **i-k,** Representative spatial visualization from IP-SNSCC case 336. **i,** Whole-section cell-coordinate map showing macrophage localization relative to the tumor reference compartment, with macrophages colored by distance-defined zones; the boxed region indicates the ROI shown in panels j and k. **j,** Magnified cell-coordinate map of the boxed ROI, with macrophages colored by mean IDO1 intensity. Background non-macrophage cells are shown in light gray and tumor/reference cells in darker gray. **k,** Matched raw PhenoCycler image from the same ROI showing DAPI, CD68 and IDO1. For spatial analyses, the reference compartment was defined as tumor in IP-SNSCC and epithelium in IP. The representative visualization in panels i-k illustrates the spatial pattern quantified across the cohort in panels a-h; quantitative conclusions were derived from the full cell-coordinate analysis rather than the selected ROI alone. Together, these analyses show that macrophages in IP-SNSCC are redistributed toward the lesional interface and exhibit nonuniform, zone-dependent IDO1 expression. Scale bars, as indicated.

### IDO1-high macrophages are enriched in IP-SNSCC

Given the interface-proximal localization of macrophages in IP-SNSCC, we next interrogated whether this compartment also exhibited an immunoregulatory phenotype. Indoleamine 2,3-dioxygenase 1 (IDO1) is a tryptophan-catabolizing enzyme expressed by activated myeloid cells and other components of the tumor microenvironment that functions as a metabolic immune checkpoint. IDO1 contributes to suppression of effector T-cell responses through local depletion of tryptophan and accumulation of immunosuppressive kynurenine metabolites. IDO1+ myeloid cells have been implicated in immune evasion across multiple solid tumors, including head and neck squamous cell carcinoma (HNSCC). To determine whether IP-SNSCC lesions harbor localized IDO1-enriched immunoregulatory niches at the tumor interface, we assessed macrophage IDO1 expression as a function of distance from the lesional compartment. Mean macrophage IDO1 expression was higher in IP-SNSCC than in IP (**Fig. 4e**), and the fraction of IDO1-high macrophages was also increased (median, 21.0% versus 4.2%), representing an approximately fivefold difference between groups (**Fig. 4f**). Higher macrophage IDO1 enrichment in IP-SNSCC was not dependent on the within-specimen all-cell Q75 threshold definition, a specimen-specific cutoff calculated as the 75th percentile of IDO1 intensity across all cells within each individual sample (**Fig. S5a-d**). Although inter-sample heterogeneity was evident in both cohorts, one benign IP specimen (IP-385) displayed an IDO1-high macrophage fraction comparable to that observed in IP-SNSCC, highlighting biological variability across lesions.

### Macrophage IDO1 expression is spatially patterned in IP-SNSCC

As macrophages were stratified into core (0-50 µm), margin (50-200 µm) and far (>200 µm) zones, IDO1 intensity distributions were compared across these regions. In IP-SNSCC, median macrophage IDO1 expression was highest in interface-adjacent zones, particularly the margin compartment (**Fig. 4g**). This spatial pattern was also evident at the individual-sample level, where IP-SNSCC specimens showing consistently higher IDO1 expression in the margin and far regions, whereas IP cases exhibited lower and more heterogeneous zonal values (**Fig. 4h**). Representative spatial maps from IP-SNSCC 336 further illustrate macrophage zone assignment across the whole section, IDO1-enriched macrophages within an interface-adjacent ROI, and the corresponding raw PhenoCycler ROI (Fig. 4i-k). Collectively, these data indicate that macrophage IDO1 expression in IP-SNSCC is spatially structured rather than uniformly distributed, with enrichment in interface-associated regions.

### The chemoattractant CXCL14 is upregulated in IP-SNSCC tumor cells

To uncover factors associated with immune cell recruitment, we analyzed genes upregulated in IP-SNSCC tumors relative to benign sinonasal papillomas. Among the top 25 genes, the conserved chemokine *CXCL14* emerged as the most highly upregulated chemokine in IP-SNSCC cells relative to IP (**Fig. 5a**, adjusted P = 8.14 × 10^-117^), implicating it as a potential mediator of immune cell composition within the IP-SNSCC microenvironment. Although *CXCL14* expression was also detected in fibroblast populations in both IP and IP-SNSCC specimens, tumor cells in IP-SNSCC exhibited significantly higher *CXCL14* expression compared to benign papilloma cells (**Fig. 5b**). In contrast, no notable differences were observed in the expression of CXC chemokines (e.g. *CXCL10*, *CXCL11*, *CXCL12*, *CXCL13*), CC chemokines (e.g. *CCL5*, *CCL19*, *CCL21*) or chemokine receptors (e.g. *CXCR3*, *CXCR5*, *CCR5*, *CCR7*) (**Fig. 5b**). To validate these transcriptomic findings in tissue, we performed RNAscope for *CXCL14* on representative IP and IP-SNSCC specimens. Consistent with the single-cell data, *CXCL14* signal was sparse to minimal in IP epithelium, whereas IP-SNSCC showed detectable punctate staining within tumor cells (**Fig. 5c, d**). Higher-magnification views highlighted abundant tumor-cell-associated *CXCL14* transcripts in IP-SNSCC, supporting epithelial/tumor-cell upregulation *in situ*. Together, these data identify *CXCL14* as a tumor-associated chemokine enriched in IP-SNSCC and nominate it as a candidate mediator of immune cell recruitment within the malignant microenvironment.

**Figure 5.**
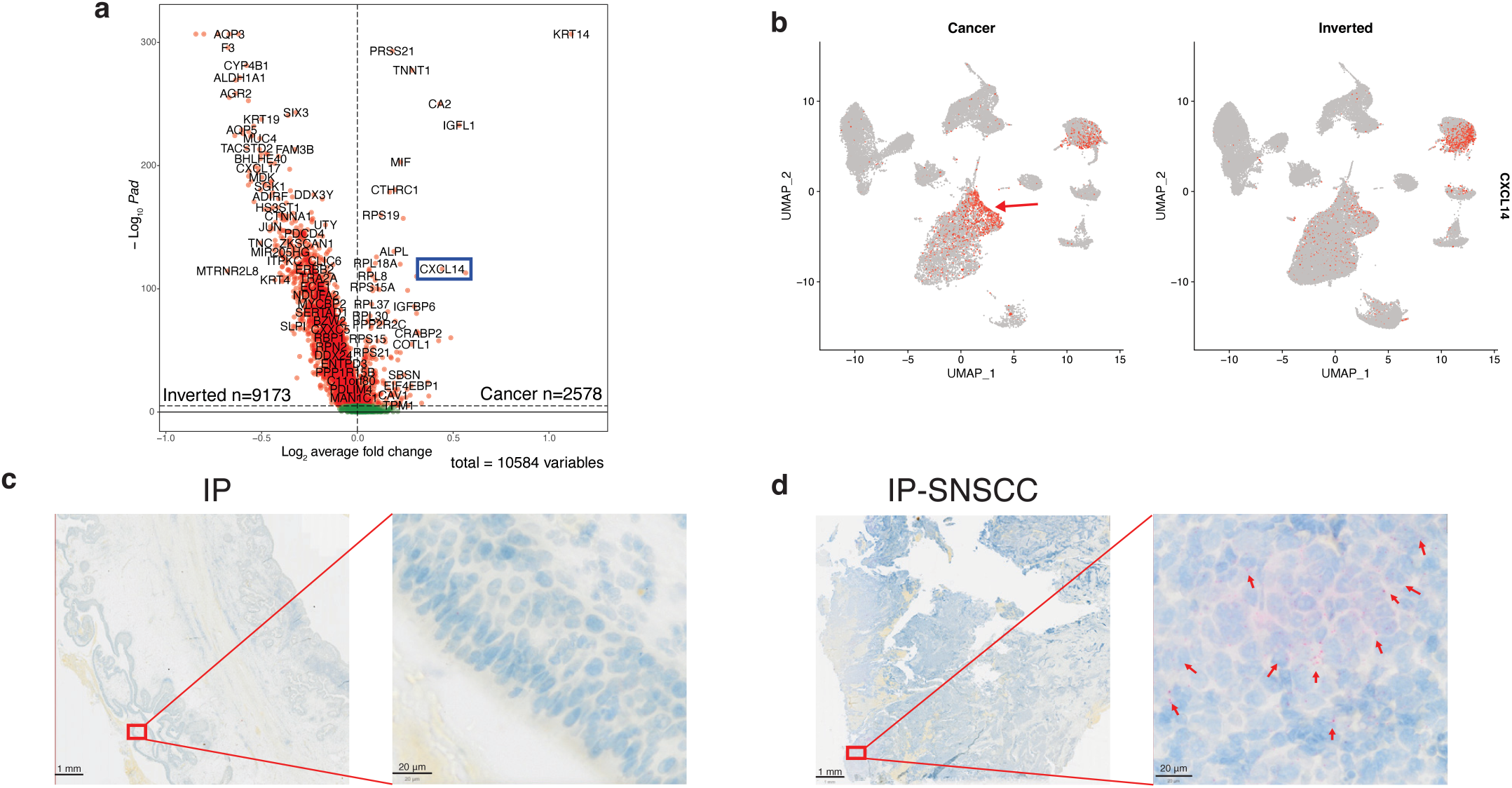
*CXCL14* is upregulated in IP-SNSCC tumor cells. a, Volcano plot showing differential gene expression in tumor cells from IP and IP-SNSCC specimens. *CXCL14* emerged as a prominently upregulated immune-related transcript in IP-SNSCC tumor cells relative to IP. **b,** Split UMAP feature plots showing *CXCL14* expression across cells from IP-SNSCC (“Cancer”) and IP (“Inverted”) specimens. In IP-SNSCC, *CXCL14* expression is concentrated in a tumor cell cluster (red arrow), whereas in IP, *CXCL14* signal is comparatively sparse. **c,d,** Representative chromogenic RNAscope images of *CXCL14* in IP (c) and IP-SNSCC (d) tissue sections. Low-magnification views are shown at left and higher-magnification insets at right. Red arrows indicate punctate *CXCL14* signal in tumor cells. Scale bars, 1 mm (low magnification) and 20 μm (high magnification).

### CXCL14 induces immunosuppressive myeloid cell phenotype

CXCL14 has been shown to directly stimulate fibroblast proliferation and migration in an autocrine manner, promoting tumor growth in prostate cancer^20^, and to mediate epithelial-to-mesenchymal transition and metastasis in a breast cancer model^21^. To determine whether CXCL14 expression by IP-SNSCC cells drives tumor growth through a similar autocrine mechanism, we treated human SNSCC-derived cell lines (NC1 and NC6), IP-SNSCC-derived cells (NC4), and a non-sinonasal SCC cell line established from oral cavity cancer (UMSCC1), with recombinant human CXCL14 (rhCXCL14) (**Fig. S6a-f**). Notably, NC1 and NC4 cells exhibited basal CXCL14 expression (**Fig. S6a**), supporting their suitability for evaluating CXCL14-mediated effects. rhCXCL14 treatment had minimal impact on proliferation in NC1 and NC6 cells (**Fig. S6b, e**), and only a modest, biologically negligible increase in NC4 cells (**Fig. S6c, d**), suggesting that direct stimulation of tumor growth is unlikely to represent the dominant mechanism of CXCL14-mediated progression. CXCL14 has also been reported to act as a potent paracrine mediator, promoting recruitment and activation of monocytes, alternatively activated macrophages, and lymphocytes^22–24^. As such, we tested whether rhCXCL14 alters immune cell activation by differentiating patient-derived monocytes and activating papilloma-infiltrating lymphocytes (PILs) or IP-SNSCC tumor-infiltrating lymphocytes (TILs) (**Fig. 6a**). CD206 expression, a well-described marker of alternatively activated or immunosuppressive “M2” macrophages enriched in our IP-SNSCC scRNA-seq tumor microenvironment dataset, was assessed by flow cytometry (**Fig. 6b**). As expected, IL-4 primed macrophages (used as a positive control) showed increased CD206 expression compared to vehicle (media only) control (**Fig. 6c**). Notably, rhCXCL14 treatment similarly induced CD206 expression in human monocyte-derived macrophages (**Fig. 6c**). In contrast, rhCXCL14 had no significant effect on activation of CD8^+^ PIL or TIL (**Fig. 6d, e**). Collectively, these findings indicate that CXCL14 may promote IP-SNSCC progression by shaping an immunosuppressive tumor immune microenvironment rather than directly modulating cytotoxic T cell activity.

**Figure 6.**
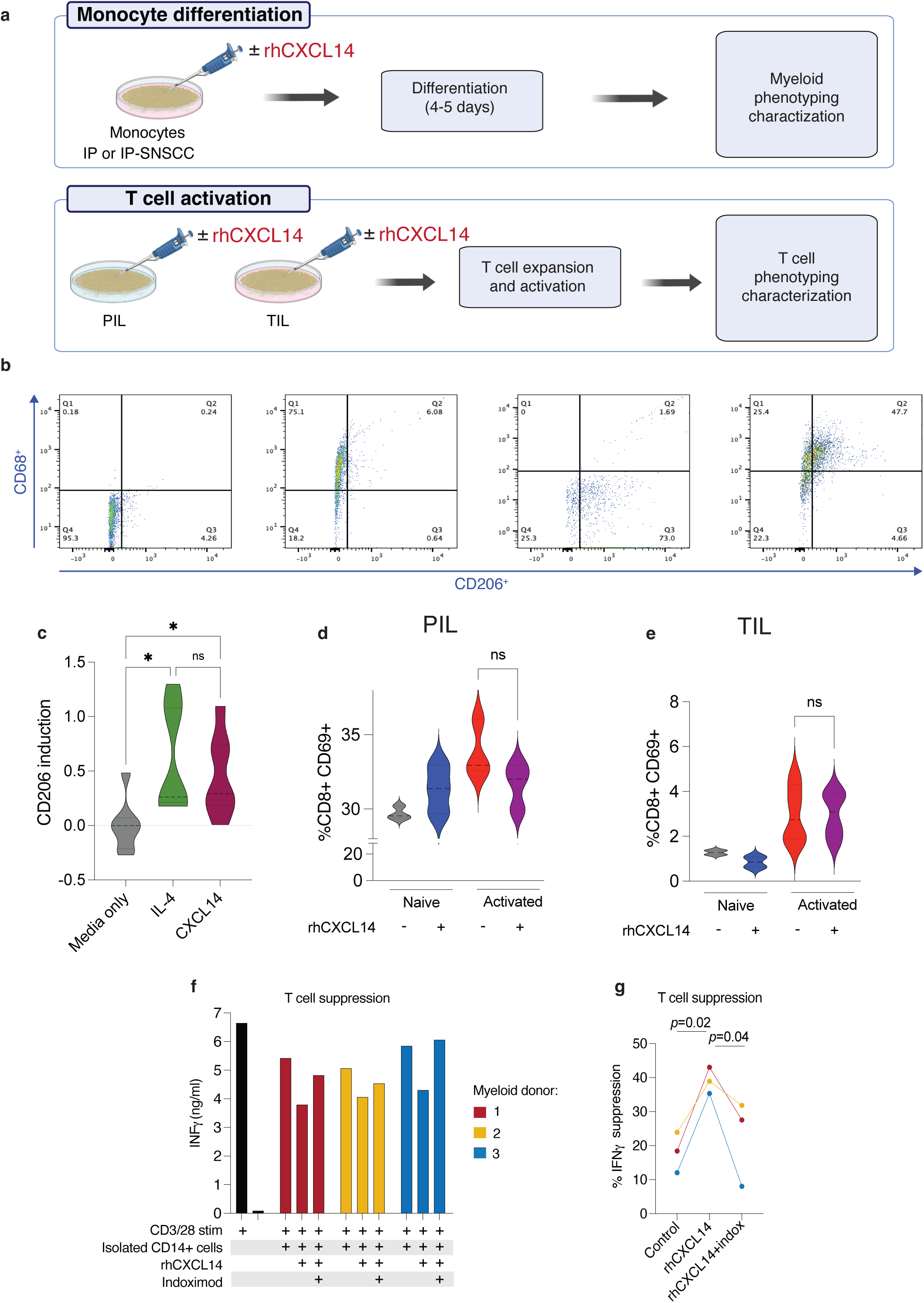
CXCL14 promotes an immunoregulatory myeloid phenotype and modulates T-cell activation. a, Schematic of the experimental design. Primary monocytes isolated from patients with IP or IP-SNSCC were differentiated in the presence or absence of recombinant human CXCL14 (rhCXCL14) for 4-5 d and then analyzed by myeloid phenotyping. In parallel, papilloma-infiltrating lymphocytes (PIL) from IP or tumor-infiltrating lymphocytes (TIL) from IP-SNSCC were expanded and activated in the presence or absence of rhCXCL14 and then analyzed by T-cell phenotyping. **b,** Representative flow-cytometry plots showing gating for unstained control, CD68^+^ cells, CD206^+^ cells and CD68^+^CD206^+^ macrophages. **c,** CD206 induction in monocyte-derived macrophages following treatment with media alone, IL-4 or rhCXCL14. **d,e,** Percentage of CD8^+^CD69^+^ cells among PIL (**d**) and TIL (**e**) under naïve and activated conditions in the absence or presence of rhCXCL14. **f,** IFNγ production by anti-CD3/CD28-stimulated T cells co-cultured with CD14+ myeloid cells from three independent donors under control, rhCXCL14, and rhCXCL14+Indoximod conditions **g,** Percentage suppression of T-cell IFNγ production relative to control for each donor. P values are indicated (p=0.02 for rhCXCL14 versus control; p=0.04 for rhCXCL14+Indoximod versus rhCXCL14). Statistical comparisons were performed using two-sided Kruskal-Wallis tests with Dunn’s multiple-comparisons correction for **c** and two-sided Mann-Whitney U tests for **d,e**; Two-way ANOVA with Tukey’s multiple comparisons test for panel **g**; ns, not significant.

### CXCL14 -conditioned myeloid cells suppresses T-cell function through an IDO-associated mechanism

Having established that CXCL14 promotes an immunoregulatory myeloid state *in vitro*, we next asked whether CXCL14-conditioned CD14+ myeloid cells suppress T-cell function, a previously unrecognized function of CXCL14. CD14⁺ monocytes from three independent healthy donors were treated with vehicle or rhCXCL14 for 24 hours and subsequently co-cultured with anti-CD3/CD28-stimulated T cells from a healthy individual. IFNγ production was measured as a readout of T-cell effector function (**Fig. 6f**). Relative to vehicle, rhCXCL14-treated CD14+ myeloid cells induced significantly greater suppression of T-cell IFNγ production (mean suppression: 39% versus 18%; p=0.02; **Fig. 6g**).

Given that CXCL14 promoted polarization of CD14+ myeloid cells toward an M2-like, tolerogenic phenotype, we hypothesized that CXCL14-conditioned myeloid cells might suppress T-cell function through an immunometabolic checkpoint program involving the IDO pathway. This hypothesis was further supported by our spatial profiling data, which identified IDO1-enriched macrophages concentrated at the tumor-stroma interface, a compartment where localized myeloid immunoregulation could directly constrain infiltrating T-cell activity. To test whether the suppressive phenotype induced by CXCL14 conditioning was associated with IDO-pathway activity, Indoximod, an IDO-pathway inhibitor, was included during the 24-hour rhCXCL14 treatment of CD14⁺ monocytes prior to co-culture with stimulated T cells. Indoximod reduced the suppressive effect of rhCXCL14-conditioned CD14⁺ myeloid cells and restored IFNγ production toward baseline levels (**Fig. 6f, g**; p=0.04 versus rhCXCL14; p=0.64 versus control). Together, these findings support a model in which CXCL14 promotes an immunoregulatory CD14⁺ myeloid state that suppresses T-cell effector activity through an IDO-pathway-associated mechanism, defining a previously unrecognized immunosuppressive role for CXCL14.

### Whole exome sequencing of IP and IP-SNSCC tumors reveals stop-gain mutation or copy number variant loss in CDKN2A as defining genomic features of IP-SNSCC transformation

To define mutational profiles across papilloma-to-carcinoma spectrum, we performed WES on tumors obtained from sinonasal papilloma and IP-SNSCC patients. In parallel, we inferred large- scale chromosomal copy-number alterations from the scRNA-seq dataset using InferCNV. Analysis of the scRNA-seq data revealed tumor cell-specific gain of chromosome 8 and loss of chromosome 5 (**Fig. S7**), consistent with copy-number alterations observed in the WES dataset (**Fig. 7a**). Across all tumors, missense mutations constituted the predominant variant type, and C>T substitutions represented the most common single-nucleotide change (**Fig. 7b**). Consistent with prior reports^7,25^, *EGFR* exon 20 mutations were recurrently observed in the cohort (**Fig. 7b**). Other frequently mutated genes included *FAT1*, *TTN*, *CKN3A*, *USP9X*, *TP53*, *CTSE*, *CSPG4*, *CLCN3*, and *ALMS1* (**Fig. 7b**). At the individual tumor level, two IP-SNSCC samples (p379 and p403) were positive for low-risk HPV by *in situ* hybridization. Missense mutations in *TP53* were detected in IP with severe dysplasia and in 2 of the 4 IP-SNSCC tumors (p168 and p264). In contrast, stop-gain mutation or copy-number variant loss of *CDKN2A* was identified in 3 of 4 IP-SNSCC specimens (**Fig. 7c**) and in none of the IP cases, aligning with the observations reported by Udager and colleagues^10^. These findings support *TP53* and *CDKN2A* alterations as defining genomic features associated with malignant transformation from IP to IP-SNSCC.

**Figure 7.**
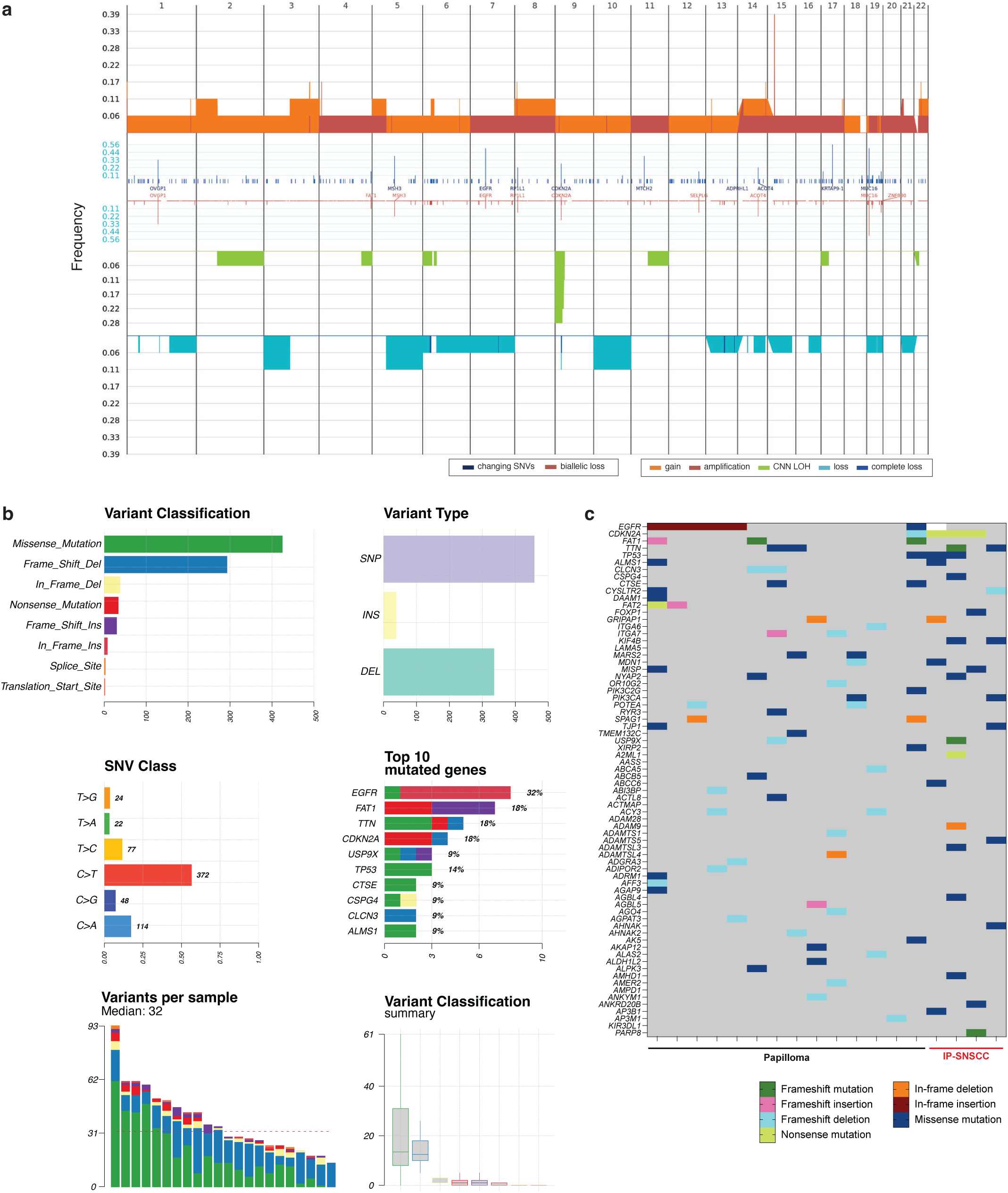
Whole-exome sequencing of IP and IP-SNSCC tumors identifies recurrent CDKN2A alterations associated with malignant transformation. a, Cohort-level summary of copy-number and sequence alterations, showing the fraction of samples with gains, amplifications, copy-neutral loss of heterozygosity (CNN-LOH), losses, complete losses, coding single-nucleotide variants (SNVs) and biallelic loss across the genome. Frequently altered genes are annotated. **b,** Summary of somatic variant features across the cohort, including variant classification, variant type, single-nucleotide substitution class, number of variants per sample, variant-classification summary and the top 10 altered genes. **c,** Oncoplot showing recurrently altered genes across all profiled tumors, including papilloma and IP-SNSCC specimens. Columns represent individual samples and rows represent genes; colors denote alteration type, including missense mutation, nonsense mutation, splice-site mutation, in-frame insertion/deletion and frameshift insertion/deletion.

## Discussion

In this study, we generated the first integrated single-cell transcriptomic and spatial proteomic map of malignant transformation from sinonasal inverted papilloma (IP) to IP-associated squamous cell carcinoma (IP-SNSCC). This multimodal analysis revealed progression as a coordinated process of epithelial, genomic and immune remodeling, implicating respiratory basal-like precursor states and recurrent *CDKN2A* loss and *TP53* alterations together with the emergence of a spatially restricted immunosuppressive myeloid niche at the tumor-stroma interface. Within this niche, CXCL14-expressing tumor cells were spatially associated with macrophages and IDO1-enriched myeloid cells at the invasive front, distinguishing IP-SNSCC from the more diffuse immune architecture of benign IP. Functional studies demonstrated that CXCL14-conditioned CD14⁺ myeloid cells suppress T-cell IFNγ production. This effect was attenuated by inclusion of Indoximod during CXCL14 conditioning prior to T-cell co-culture, supporting an IDO-pathway-associated mechanism underlying myeloid-mediated T-cell suppression. Together, these findings define a previously unrecognized CXCL14-IDO immunoregulatory program at the invasive front and suggest that malignant progression in IP-SNSCC is shaped not only by epithelial transformation and tumor suppressor loss, but also by spatially localized immune restraint. By resolving context-dependent immune mechanisms that are difficult to infer from bulk or dissociated analyses, our work positions the CXCL14-IDO pathway as a candidate therapeutic vulnerability worthy of systematic evaluation in other immune-rich but functionally restrained cancers.

Beyond transcript abundance, our spatial analyses indicate that the transition from IP to IP-SNSCC is accompanied by a shift in immune positioning toward the lesional compartment: multiple distance-based and proximity metrics (including immune-to-lesion median distance and related proximity indices) consistently showed increased immune cell proximity in IP-SNSCC, independent of total cellularity and overall immune fractions. Similar distance-to-tumor and compartmentalized infiltration measurements have been used in multiplexed tissue profiling studies to distinguish “immune-infiltrated” versus “immune-excluded” architectures and link spatial proximity to functional immune states^34,35^.

We identified enrichment of myeloid cells, particularly alternatively activated or immunosuppressive macrophages, as a defining feature of the IP-SNSCC tumor microenvironment. Prior studies have demonstrated that CXCL14 acts as a chemoattractant for multiple immune cell types within tumors, including dendritic cells^36^, natural killer cells^37^, CD8+ T cells^24^, monocytes^20^, and alternatively activated (M2) macrophages^38^. Its role as a tumor-suppressive or tumor-promoting chemokine is highly context-dependent, influenced by cell-type-specific expression patterns and the composition of the recruited immune populations. Consistent with this, CXCL14 restoration has been shown to enhances anti-tumor T cell immunity in HPV-driven head and neck cancer, underscoring that its effects vary with tumor type, antigenicity, and the infiltrated immune contexture^30^.

Notably, beyond its role in immune cell recruitment, CXCL14-dependent M2 macrophage polarization, including enrichment of CD206^+^ macrophages, has been described in the osteogenic tumor microenvironment^38^, sepsis^23^, and adipose tissue^22^, with recent mechanistic studies implicating NF-κB mediated signaling in cancer-associated contexts^22,32^. Our study expands the biological scope of CXCL14 in cancer by defining a previously unrecognized immunomodulatory role in IP-SNSCC, linking tumor-derived CXCL14 upregulation to the accumulation of alternatively activated macrophage states within a spatially organized, localized immunosuppressive microenvironment. Moreover, our data demonstrate that CXCL14 directly induces alternatively activated or immunosuppressive macrophage phenotype. Although several receptors have been proposed to mediate CXCL14 signaling, including CXCR4^39,40^, ACKR2^41,42^, and MRGPRX2^43^, the definitive receptor responsible for CXCL14-mediated effects remains unknown and requires further elucidation.

In the spatial domain, macrophages mirrored the global immune proximity pattern, with relative enrichment at the lesion-stroma interface in IP-SNSCC. Interface-localized macrophage niches have been reported across tumor types using multiplexed platforms, where their positioning and local neighborhoods are linked to immunoregulatory programs at invasive fronts and tumor borders^13,44^. Notably, our PhenoCycler data indicate that IDO1 expression is spatially restricted and enriched in interface-adjacent myeloid regions in IP-SNSCC, consistent with the concept of a localized “immunoregulatory niche” at the tumor border. Prior spatial studies have documented boundary-associated IDO1 in interferon/inflammation-rich transition zones, and extensive evidence supports IDO1 as an interferon-inducible metabolic immune checkpoint that can suppress effector T/NK cell function and promote regulatory programs^45,46^.

However, spatial data alone could not determine whether CXCL14 merely co-localized with these interface-associated immunoregulatory features or actively contributed to their establishment. Our functional assays provide mechanistic support linking CXCL14 upregulation to an IDO-pathway-associated program of T-cell suppression. Although our spatial analyses initially identified a correlation between interface-localized IDO1-high macrophages and CXCL14-rich IP-SNSCC lesions, the co-culture data extend this observation by showing that rhCXCL14 treatment of CD14+ myeloid cells promotes an immunoregulatory state that enhances suppression of T-cell IFNγ production. Restoration of IFNγ production by Indoximod further supports the involvement of the IDO pathway in this suppressive effect.

These findings may help explain a central feature of IP-SNSCC: despite an immune-rich microenvironment, anti-tumor immune function appears constrained rather than fully productive. We propose that CXCL14 contributes to this state by promoting a localized myeloid suppressive program associated with IDO-pathway activity, thereby limiting effective T-cell responses at the tumor interface. More broadly, this model suggests that immune cell abundance alone may not fully capture the functional immune state of these tumors and could therefore complicate biomarker-based stratification for immunotherapy. Together, these data identify a previously unrecognized immunoregulatory mechanism associated with the CXCL14-IDO pathway in IP- SNSCC and support further investigation of this pathway in other tumor contexts characterized by immune-rich but functionally restrained microenvironments.

The cellular origin of sinonasal papillomas has yet to be defined. Here, we identify basal cells of the respiratory epithelium^47^ as likely precursors for IP development. As epithelial cells transition from normal respiratory epithelium to papillomatous tumors, we observe loss of cilia with concurrent upregulation of basal epithelial cell markers. Consistent with this, impaired ciliogenesis is a recognized hallmark of tumorigenesis across multiple cancer types^48,49^. Pseudotime analysis further delineated two distinct trajectories, one characterized by aberrant proliferation consistent with papillomatous growth and transformation, and the second marked by altered extracellular matrix programs linked to cancer progression and metastasis^50^. In parallel, prior targeted next-generation sequencing studies of IP and IP-SNSCC tumors have identified frequent enrichment of *EGFR* exon 20 activating mutations and *CDKN2A* alterations in IP-SNSCC tumors^7,9,10^. We extend these observations using high-throughput whole-exome sequencing, demonstrating that truncating *CDKN2A* alterations and *TP53* mutations are defining genomic features of malignant transformation to IP-SNSCC.

Taken together, these findings provide the first comprehensive molecular characterization of the IP-SNSCC tumor immune microenvironment and identify CXCL14 as a potential novel immunomodulatory driver of malignant transformation from IP to IP-SNSCC through a localized myeloid immunoregulatory program associated with IDO-pathway activity (**Fig. S8**). In this framework, enrichment of immunosuppressive myeloid populations and interface-associated IDO1 expression may help explain how immune-rich lesions remain functionally constrained, thereby identifying CXCL14-associated myeloid immunoregulation as a candidate therapeutic vulnerability in sinonasal squamous cell carcinoma. Collectively, these results also identify a newly discovered immunosuppressive function of CXCL14 which was previously unrecognized.

### Study considerations and limitations

The spatial analyses were performed on two-dimensional tissue sections, and architectural differences between papillary IP lesions and more solid IP-SNSCC tumor nests can influence distance-based measurements; nevertheless, distance-to-tumor and compartment-specific infiltration metrics are widely used in multiplex imaging studies and remain informative when interpreted alongside morphology^34,51^. In addition, the cohort size for high-plex spatial profiling was limited, restricting formal inference and motivating validation in larger, clinically annotated datasets. Furthermore, although our co-culture experiments provide functional support for a CXCL14-linked myeloid suppressive mechanism associated with IDO-pathway activity, they do not establish direct molecular regulation of IDO1 by CXCL14 or fully recapitulate the complexity of the intact tumor microenvironment. Additional mechanistic and *in vivo* studies will be needed to determine how CXCL14 shapes myeloid cell states within IP-SNSCC, to define the precise relationship between CXCL14 and IDO-pathway activation, and to evaluate whether this immunoregulatory mechanism can be therapeutically exploited.

## Materials and Methods

### Patient Samples

Patients were prospectively enrolled in this study at Johns Hopkins. All participants provided written informed consent prior to inclusion in the study. Studies were performed in accordance with the Institutional Review Boards at the National Institutes of Health and Johns Hopkins. The scRNA-seq cohort initially included 20 patients (15 benign sinonasal papillomas and 5 IP-SNSCC) and 22 samples, with subject 289 contributing three scRNA-seq samples (IP-2, IP-3, and IP-4) from a multifocal IP, sampled from the inferior turbinate, middle turbinate, and septal attachments. Following scRNA-seq quality assessment, 19 patients (15 benign sinonasal papillomas and 4 IP-SNSCC) and 21 samples were retained for downstream analysis, as the sample from subject 336 was excluded based on QC metrics. Of these 19 patients, 17 also underwent WES, whereas one benign papilloma patient (IP-1) and one IP-SNSCC patient (IP-SNSCC-1) were analyzed only by scRNA-seq. WES was additionally performed in one IP-SNSCC patient outside the scRNA-seq cohort (subject 379), bringing the total WES cohort to 18 patients (14 benign sinonasal papillomas and 4 IP-SNSCC) and 36 DNA samples (18 tumor-normal pairs). Demographic and clinical variables collected for the cohort included age at diagnosis, sex, race/ethnicity, and smoking history.

### Mammalian cell lines and culture

NC1, NC4, NC6 sinonasal squamous cell carcinoma-derived cell lines were obtained from Mario Hermsen (Hospital Universitario Central de Asturias, Oviedo, Asturias, Spain) and have been previously described (Garcia-Inclan et al. 2014). Cells were maintained in minimal essential medium (MEM) culture, supplemented with 10% FBS, 100 U/ml penicillin, 200 μg/ml streptomycin, 2 mM L-Glutamine and non-essential amino acids 100 μM and incubated in 5% C0_2_ at 37°C. UMSCC-1 were obtained from Thomas Carey (University of Michigan, Michigan, USA) and maintained in MEM with 10% fetal bovine serum, penicillin/streptomycin, l-glutamine, and nonessential amino acids. All cells were harvested for experimental use at low passage number and serially tested for mycoplasma every 3 months.

### Tumor dissociation

Patient tissue was diced into 1 mm^3^ pieces and dissociated into a single cell suspension using the Miltenyi Biotec Tumor Dissociation Kit, human (130-095-929). The single-cell suspension was centrifuged at 300 × g for 5 min and resuspended in 0.5 mL of Red Blood Cell Lysis Buffer 10x (BioLegend 420301) and 4.5 mL sterile water. After 1 min, the reaction was quenched with 20 mL RPMI 1640. The cells were spun down, and all remaining liquid was aspirated off. The pellet was resuspended in 1% FSB (0.5 g BSA in 50 mL PBS, sterile filtered). The suspension was run through a 35 μm strainer and counted using standard AO/PI staining.

### scRNA-sequencing and processing

scRNA-seq of isolated tumor cells was performed using the 10X Single Cell 5′ Gene Expression Kit (10X Genomics, Pleasanton, CA, USA). Cells were captured in droplets at a targeted cell recovery of 5000 to 10,000 cells per lane and scRNA library generation was performed. After cell barcoding and reverse transcription, emulsions were broken and cDNA purified using Dynabeads. Purified cDNA was then amplified by at least 10 polymerase chain reaction (PCR) cycles for 5′ gene expression. Amplified cDNA was then used for 5′ gene expression library construction. Sequencing was performed using an Illumina NovaSeq 6000 with 310 million reads per sample and a sequencing configuration of 26 x 8 x 98 (UMI x Index x Transcript read). The Cell Ranger 3.0.2. pipeline software (10X Genomics, California) was used to align reads and generate expression matrices for downstream analyses. Automatically called cells were further filtered to retain high-quality droplets with captured cells using thresholds of >500 total UMI counts, >250 detected features, and <25% mitochondrial transcripts.

### Single-cell data integration and differential expression

Downstream single-cell analyses were performed in Seurat v4.3.0^52^. Data were normalized, variable features were identified, gene expression values were scaled, and principal component analysis was performed. Uniform Manifold Approximation and Projection (UMAP) was used for dimensionality reduction, and shared nearest-neighbor graphs followed by Leiden clustering were used to define cell populations. Batch-effect correction across samples was performed using Harmony^53^. Clusters were annotated using a combination of canonical marker genes and differential expression comparing each cluster with all remaining cells in the dataset. Differential expression analyses were performed using the Mann-Whitney U test.

### Gene set enrichment analysis

Gene set enrichment analysis (GSEA) was performed using fgsea v1.24.0^54^ with gene sets obtained from the Molecular Signatures Database (MSigDB)^55^. Features were ranked by - log(Pvalue) × sign(fold change).

### Cluster abundance analysis

Percentages of cells by cluster and sample were calculated as the proportion of cells in a given cluster relative to the total number of cells in each sample. Statistical comparisons of cluster proportions were performed using t tests.

### RNA velocity and pseudotime analysis

For RNA velocity analyses, reads were assigned as spliced or unspliced using velocyto v0.17.17^56^, and velocity embeddings were generated using scVelo v0.2.5^57^. Pseudotime trajectories were inferred using Slingshot^58^. A generalized additive model was used to identify genes varying significantly over pseudotime, and z-scored expression values were used to visualize the most significantly dynamic genes along each trajectory.

### Intercellular signaling analysis

Intercellular signaling analyses were performed using Domino^59^. UCell^60^ was used to calculate transcription factor activation scores for each cell using transcription factor target gene sets from MSigDB^61^. CellPhoneDB v2^62^ was used to identify candidate ligand-receptor pairs, and the NicheNet protein-protein interaction network was used to identify receptors upstream of transcription factors in known signaling networks. Signaling scores between cell clusters were calculated as the product of z-scored ligand expression, receptor expression, and transcription factor activation scores. For each transcription factor, the highest inter-cluster score was retained, and summed scores were used to quantify overall signaling between clusters. ComplexHeatmap v2.14.0^63^ and circlize v0.4.15^64^ were used for visualization.

### Deriving cell-based and spatial features

Cell-based and spatial feature derivation have been previously described^65^. Briefly, for each sample, a 39-length vector was computed with the average biomarker pixel intensity across the sample image for each biomarker and with the average biomarker pixel intensity within the cell segmentation masks for each biomarker. A 16-length vector was then computed with the fraction of each cell type present in the sample. This vector summed to 1.

### Neighborhood matrix

For each sample, we computed a “neighborhood matrix,” M, which is a matrix with shape (number of cell types) × (number of cell types), and each row sums to 1. Here, M is a 16 × 16 matrix. Element m_ij is the fraction of cell type j within the k-nearest neighbors of cell type i, averaged across all cells of type i in the sample. In this work, we use k = 10, and nearest neighbors are computed using the centroid coordinates of each cell.

### PhenoCycler-Fusion system

Multiplex immunofluorescence imaging was performed on six tissue specimens, including three inverted papilloma (IP) and three IP-associated sinonasal squamous cell carcinoma (IP-SNSCC) samples, using the PhenoCycler-Fusion platform (Akoya Biosciences). PhenoCycler 25-plex panel was designed as previously described^66^. Preselected antibody clones were conjugated to DNA oligonucleotide barcodes using a conjugation kit from Akoya Biosciences (SKU 7000009) according to the manufacturer’s instructions, with each barcode corresponding to a fluorophore reporter molecule that could be visualized on the Alexa Fluor 647 (AF647), ATTO550, or Alexa Fluor 750 (AF750). FFPE samples were sectioned onto microscope slides, baked at 60 °C overnight in an incubator, deparaffinized with xylenes, rehydrated with alcohol, and washed twice with Milli-Q ultrapure water. Antigen retrieval was then performed using AR9 buffer (SKU AR9001KT, Akoya Biosciences). Custom-conjugated antibodies were prepared according to the manufacturer’s instructions (SKU 7000017, Akoya Biosciences) and incubated in post-staining fixing solution (SKU 232107, Akoya Biosciences). Images were obtained using the PhenoCycler-Fusion 2.0 device.

### Marker quality control

Marker quality was assessed for each channel using a predefined score- and interpretation-based framework. Channels with scores <2 or interpreted as non-specific binding, no signal, or weak signal were excluded from analysis, whereas channels with scores ≥2 and interpreted as good staining or saturated were retained. Markers flagged for exclusion were removed from phenotyping analyses. CD45RO and PD-1 were excluded from both cohorts. In the IP cohort, PD-L1 and Pan-Cytokeratin were additionally excluded because of out-of-focus artifacts.

### Cell segmentation, phenotype classification, and compartment assignment

Cell segmentation and marker quantification were performed in QuPath (v0.6.0) using the StarDist extension (v0.6.0). Segmentation was based on the DAPI channel using a requested pixel size of 0.5 µm, with nucleus expansion of 2-3 µm to capture cytoplasmic signal. Mean marker intensities were quantified for each segmented cell across all channels. Cell phenotypes were assigned in QuPath using a hierarchical gating strategy based on marker intensity thresholds. Thresholds were determined interactively using QuPath live preview feature, with values set based on visual inspection of positive and negative control regions and validated across multiple representative specimens. For immune phenotypes, CD45 positivity was established as a prerequisite gate, followed by sequential classification into specific immune subtypes based on additional marker expression. Harmonized phenotype classes across specimens included epithelial cells, macrophages, cytotoxic T cells, helper T cells, regulatory T cells, B cells, smooth muscle cells, and other/unclassified cells. Phenotype assignments were validated by visual inspection to ensure consistency across specimens. For downstream spatial analyses, phenotype labels were grouped into broader analytic compartments using rule-based mapping. The Immune compartment included T-cell, B-cell, macrophage, and other immune-related phenotypes, whereas the Stromal compartment included smooth muscle and other stromal-associated phenotypes. Epithelial-like phenotypes (defined by expression of cytokeratin, CK, E-cadherin, or Pan-Cytokeratin) were assigned in a disease-context-specific manner to the cohort-specific epithelial-associated compartment used for spatial analysis, here termed the *lesional compartment*; this was defined as the Epithelial compartment in IP specimens and the Tumor compartment in IP-SNSCC specimens. Cells that did not meet classification criteria were labeled as Unknown.

### Spatial distance analysis

Per-cell QuPath exports comprising centroid coordinates, phenotype labels, compartment labels, and mean marker intensities were imported into Python for downstream analysis. When QuPath exported coordinates in microns, values were used directly; when coordinates were in pixels, values were converted using the imaging resolution of 0.5 µm/pixel. For cohort-level spatial analyses, the reference compartment was defined as the Epithelial compartment in IP and the Tumor compartment in IP-SNSCC. Euclidean distance from each immune cell to the nearest reference-compartment cell was computed using KDTree nearest-neighbor search in SciPy (v1.12.0). Immune cells were stratified into distance bins of 0-50 µm, 50-100 µm, 100-200 µm, and >200 µm. The infiltration ratio was defined as the fraction of immune cells within 50 µm of the nearest reference-compartment cell. The immune exclusion score was defined as the fraction of immune cells located at distances >200 µm divided by the fraction located within 0-200 µm. Median immune-to-reference distance was also computed for each specimen.

### Macrophage proximity analysis

Macrophages were identified from QuPath-assigned phenotype labels. For each macrophage, the minimum Euclidean distance to the nearest reference-compartment cell was computed using KDTree nearest-neighbor search in SciPy (v1.12.0). Macrophages were then stratified into three spatial zones according to this distance: core (0-50 µm), margin (50-200 µm), and far (>200 µm). Summary metrics included macrophage infiltration ratio, defined as the fraction of macrophages within 50 µm of the reference compartment; median macrophage-to-reference distance; zone fractions; and a macrophage infiltration score defined as the ratio of the core fraction to the far fraction.

### Macrophage IDO1 analysis

Per-cell mean IDO1 intensity was quantified in macrophages from each specimen. Cohort-level macrophage IDO1 signal was summarized as mean and median values per specimen. To define a relative IDO1-enriched macrophage compartment, we calculated the 75th percentile of IDO1 intensity across all cells within each specimen and classified macrophages exceeding this specimen-specific all-cell threshold as IDO1-high. This threshold was used as a within-specimen enrichment metric and was not interpreted as an absolute positivity threshold. The fraction of IDO1-high macrophages was calculated for each specimen. As a sensitivity analysis, macrophage IDO1-high fractions were also recalculated using fixed study-wide and benign-reference thresholds, including pooling all-cell and macrophage-specific IP reference thresholds at the 75th and 90th percentiles. To assess spatial variation in macrophage IDO1 expression, macrophages were grouped into the same core, margin, and far zones defined in the macrophage proximity analysis, and zonal mean and median IDO1 values were computed. Within-sample two-sided Mann-Whitney U tests were used to compare core versus far macrophage IDO1 distributions.

### RNAscope

RNA in situ hybridization was performed using the RNAscope LS 2.5 Hs-CXCL14 probe (Advanced Cell Diagnostics, Bio-Techne; Cat. No. 425298), targeting human *CXCL14* mRNA. FFPE tissue sections obtained from IP and IP-SNSCC patients were processed according to the manufacturer’s instructions using the BOND RX automated stainer (Leica Biosystems). Images were acquired and analyzed on a Zeiss LSM780 confocal laser-scanner.

### PIL and TIL culture

TIL isolation and culture have been previously described (Redman JM et al. JCI 2022 PMID: 35727629) (Samaniego C et al. Head Neck 2023 PMID: 37480219). Briefly, tumors obtained from IP patients (PIL) or IP-SNSCC patients (TIL) were sectioned into 1mm^3^ fragments and cultured in media (50% AIM V media, 50% RPMI media, 25 mM HEPES buffer, 5% heat-inactivated human serum, 100 U/mL penicillin, 100 μg/mL streptomycin, 10 μg/mL gentamycin, 2 mM l-glutamine, 1.25 μg/mL amphotericin B) supplemented with 6,000 U/mL IL-2. T cells were isolated by immunomagnetic T cell negative selection using the EasySep Human T cell isolation kit (STEMCELL Technologies) according to the manufacturer’s instructions, and cryopreserved in heat-inactivated FBS with 10% DMSO.

### Peripheral blood mononuclear isolation and culture

Monocytes from three separate healthy donors were isolated from whole blood buffy coats following CPT centrifugation using the EasySep Human CD14 Positive Selection Kit II (Stemcell Technologies) per manufacturer recommendations. Fresh monocytes were suspended in media (RPMI1640 media supplemented with 10% FBS, 2 μmol/L β-ME, HEPES, nonessential amino acids, glutamine, and antibiotics) at 5x10^5^ cells/mL and exposed to recombinant human CXCL14 (100 ng/mL, R&D Systems, Cat. No. 866-CX) alone or in combination with Indoximod (50 μM, MedChemExpress; Cat. No. HY-16724), or control (volume equivalent DMSO) for 24 hours at 37°C/5%CO_2_. Cultured monocytes were then used in T cell suppression assays.

### Flow cytometry

Cells were counted by trypan blue exclusion and plated at a density of 5 x 10^5^ cells per well of a 12 well plate. PIL and TIL were activated with CD3/CD28 Dynabead (ThermoFisher Scientific) according to the manufacturer’s instructions in the presence or absence of 100 ng/mL rhCXCL14 (R&D Systems). Monocytes were cultured with 100 ng/mL rhIL-4 (R&D Systems) or rhCXCL14 for 4-5 days. Cells were washed with PBS and labeled with SYTOX blue dead cell stain (ThermoFisher Scientific) for 15 minutes at room temperature and blocked with Fc receptor binding inhibitor polyclonal antibody (eBioscience). Cells were then stained with fluorochrome-labeled antibodies, CD8, CD69, CD68, CD206 (Biolegend). Data acquisition was performed using an LSRII flow cytometer (BD Biosciences) and populations were gated to exclude debris, dead cells, and select single cells. Data analysis was performed using FlowJo (BD Biosciences).

### T cell suppression assays

T cells were isolated from PBMC from a common healthy donor via negative magnetic selection using the EasySep Human T Cell Enrichment Kit from Stemcell Technologies, per the manufacturer’s recommendations. T cells were stimulated with plate-bound anti-human CD3 (clone OKT3, 5 μg/mL) and CD28 (CD28.2, 5 μg/mL) and cultured alone or co-cultured with isolated pre-conditioned monocytes at a 2:1 myeloid-to-T cell ratio for 72 hours. IFNγ concentration in the culture supernatant was determined using an IFNγ ELISA (R&D Systems) according to the manufacturer’s recommendations. Percent IFNγ suppression for each condition was calculated as: (IFNγ concentration in CD3/28 stim condition - IFNγ concentration in experimental condition) / IFNγ concentration in CD3/28 stim condition.

### DNA Extraction

DNA was extracted from 20-30mg frozen tissue samples using the AllPrep DNA/RNA Mini Kit (Qiagen). Tissue was homogenized in 600 μL Buffer RLT Plus with β-mercaptoethanol (β-ME) using the TissueRuptor II (Qiagen). After centrifugation, the supernatant was transferred to AllPrep DNA spin columns. The column was centrifuged at ≥8000 x g for 30 seconds. Genomic DNA was purified by adding 500 μL Buffer AW1, centrifuging, discarding the flow-through, applying 500 μL Buffer AW2, and centrifuging again. Finally, 100 μL Buffer EB was added, incubated for 5 minutes, and centrifuged for DNA elution. The extracted DNA was stored at -20 °C. PBMCs were prepared for DNA extraction using the QIAamp DNA Mini and Blood Mini (QIAGEN, 51104). Genomic DNA extraction from PBMC samples was performed by GENEWIZ (Azenta Life Sciences) using standard protocols.

### Whole Exome Sequencing

Whole exome sequencing and analysis were performed as previously described ^43^. Briefly, libraries for whole exome sequencing were constructed using the Agilent SureSelect V5 post-capture beginning with a minimum of 200ng of genomic DNA and the SureSelect^XT^ Target Enrichment System for Illumina paired-end multiplexed sequencing. Paired-end sequencing was performed using the NovaSeq 6000 S4 Flow Cell with 2 × 150 cycle chemistry with an output of approximately ∼8Gb (150X) per sample. Library preparation and whole-exome sequencing from PBMC samples were performed by GENEWIZ (Azenta Life Sciences) using standard protocols.

### Somatic variant detection and annotation

Whole-exome sequencing data were processed using the nf-core/sarek workflow (v3.3.2) executed with Nextflow v23.04.4 in a reproducible Singularity-based environment. Reads were aligned to the GRCh38 human reference genome using bwa-mem2. Somatic single-nucleotide variants (SNVs) and small insertions/deletions (indels) were identified in paired tumor-normal mode using matched PBMC-derived normal DNA with GATK Mutect2 v4.4.0.0 and Strelka v2.9.10. The workflow incorporated a panel of normals and population germline resources to reduce technical artifacts and likely germline calls. Somatic variants were functionally annotated using Ensembl VEP v110. For cohort-level analyses, annotated variants were summarized by variant classification, variant type, single-nucleotide substitution class, and number of consensus variants per sample. Recurrently altered genes were identified according to alteration frequency across the cohort.

### Copy-number analysis

Copy-number profiles were computed using superFreq v1.3.2. Gene-level copy-number events were categorized as gain (3 copies), amplification (≥4 copies), copy-neutral loss of heterozygosity (CNN-LOH), loss, or complete loss. Cohort-level copy-number summaries were calculated as the fraction of individuals harboring each event per gene, and integrated visualizations were generated to display the combined frequencies of copy-number and sequence alterations across the cohort.

### Statistical analysis

Unless otherwise indicated, statistical analyses were performed using GraphPad Prism v10.2.3. Two-sided Mann-Whitney U tests were used for comparisons between two groups of biological replicates, including between-cohort comparisons of macrophage proximity and macrophage IDO1 summary metrics. Cohort-level immune spatial metrics were summarized descriptively. Within-sample core-versus-far comparisons of macrophage IDO1 distributions were performed using two-sided Mann-Whitney U tests, and exact *P* values are reported where relevant. For comparisons between two groups of technical replicates, two-tailed *t* tests were used. For comparisons involving more than two groups, two-sided Kruskal-Wallis tests with Tukey’s multiple-comparisons correction were performed. *P* < 0.05 was considered statistically significant. For scRNA-seq differential expression analyses, *P* values were adjusted for multiple testing using the Benjamini-Hochberg procedure, and genes with false discovery rate (FDR) < 0.05 were considered significant.

### Software

Single-cell analyses were performed using Seurat v4.3.0, Harmony, fgsea v1.24.0, velocyto v0.17.17, scVelo v0.2.5, Slingshot, Domino, UCell, CellPhoneDB v2, NicheNet, ComplexHeatmap v2.14.0, and circlize v0.4.15. Image analysis was performed in QuPath (v0.6.0) with the StarDist extension (v0.6.0). Downstream spatial analysis was implemented in Python using NumPy (v1.26.4), Pandas (v2.1.4), SciPy (v1.12.0), Matplotlib (v3.8.2) and Seaborn (v0.13.2). Statistical analyses were performed in GraphPad Prism v10.2.3. Figures and illustrations were assembled using Adobe Illustrator (Adobe Inc.) and BioRender.com.

## Data and code availability

The genomic data are available under controlled access due to patient privacy regulations, informed consent restrictions, and institutional ethics policies. Sequencing data generated in this study for patients with informed consent allowing data sharing have been deposited in dbGaP and can be accessed by submitting a data access request through dbGaP. Remaining sequencing data are available upon request from the corresponding author with a completed data transfer or data use agreement. Per-cell data tables and analysis scripts supporting this study will be available on GitHub, and a Zenodo DOI for the archived release will be provided upon publication.

## Acknowledgements

This study was presented at the American Head and Neck Society (AHNS) Annual Meeting at the Combined Otolaryngology Spring Meetings (COSM) in New Orleans, Louisiana, on May 14-15, 2025, where it was recognized with the Best Resident Basic Science Research Paper Award. This work utilized the computational resources of the NIH HPC Biowulf cluster (https://hpc.nih.gov).

## Author contributions

**Author contributions**: T.F.O., F.T.Z. and N.R.L. conceptualized the study. T.F.O., K.V., S.G., R.V. and A.H. performed experiments. T.F.O., F.T.Z., Y.D., Y.H., X.Y., F.C., M.N., C.C., M.P. N.W. and N.R.L. performed computational and bioinformatic analyses. T.F.O., F.T.Z., A.P.L., M.R.J., N.R.R., T.R.D., J.K., W.K., M.L., A.U., C.T.A. and N.R.L. collected clinical samples or clinical data. T.F.O., F.T.Z., K.V., S.G., Y.D., Y.H., X.Y., R.V., A.H., F.C., M.N., C.C., M.P., N.W., T.R.D., J.K., W.K., A.P.L., M.R.J., N.R.R., M.L., A.U., E.I., C.T.A. and N.R.L. interpreted the data. T.F.O., F.T.Z. and N.R.L. wrote the original draft of the manuscript. All authors reviewed, edited and approved the final manuscript.

## References

1 Thawani, R. et al. The contemporary management of cancers of the sinonasal tract in adults. CA Cancer J Clin 73, 72–112 (2023). 10.3322/caac.21752

2 Sanghvi, S. et al. Epidemiology of sinonasal squamous cell carcinoma: a comprehensive analysis of 4994 patients. Laryngoscope 124, 76–83 (2014). 10.1002/lary.24264

3 Hermans, R., De Vuysere, S. & Marchal, G. Squamous cell carcinoma of the sinonasal cavities. Semin Ultrasound CT MR 20, 150–161 (1999). 10.1016/s0887-2171(99)90016-1

4 Nicolai, P. et al. Endoscopic surgery for malignant tumors of the sinonasal tract and adjacent skull base: a 10-year experience. Am J Rhinol 22, 308–316 (2008). 10.2500/ajr.2008.22.3170

5 von Buchwald, C. & Bradley, P. J. Risks of malignancy in inverted papilloma of the nose and paranasal sinuses. Curr Opin Otolaryngol Head Neck Surg 15, 95–98 (2007). 10.1097/MOO.0b013e3280803d9b

6 Wang, Z. et al. Sinonasal Inverted Papilloma-Associated and De Novo Squamous Cell Carcinoma: A Tale of Two Cities or Not. Cancers (Basel) 14 (2022). 10.3390/cancers14215211

7 Udager, A. M. et al. Human papillomavirus (HPV) and somatic EGFR mutations are essential, mutually exclusive oncogenic mechanisms for inverted sinonasal papillomas and associated sinonasal squamous cell carcinomas. Ann Oncol 29, 466–471 (2018). 10.1093/annonc/mdx736

8 Syrjanen, K. & Syrjanen, S. Detection of human papillomavirus in sinonasal papillomas: systematic review and meta-analysis. Laryngoscope 123, 181–192 (2013). 10.1002/lary.23688

9 Udager, A. M. et al. High-Frequency Targetable EGFR Mutations in Sinonasal Squamous Cell Carcinomas Arising from Inverted Sinonasal Papilloma. Cancer Res 75, 2600–2606 (2015). 10.1158/0008-5472.CAN-15-0340

10 Brown, N. A. et al. TP53 mutations and CDKN2A mutations/deletions are highly recurrent molecular alterations in the malignant progression of sinonasal papillomas. Mod Pathol 34, 1133–1142 (2021). 10.1038/s41379-020-00716-3

11 de Visser, K. E. & Joyce, J. A. The evolving tumor microenvironment: From cancer initiation to metastatic outgrowth. Cancer Cell 41, 374–403 (2023). 10.1016/j.ccell.2023.02.016

12 Mempel, T. R., Lill, J. K. & Altenburger, L. M. How chemokines organize the tumour microenvironment. Nat Rev Cancer 24, 28–50 (2024). 10.1038/s41568-023-00635-w

13 Matusiak, M. et al. Spatially Segregated Macrophage Populations Predict Distinct Outcomes in Colon Cancer. Cancer Discov 14, 1418–1439 (2024). 10.1158/2159-8290.CD-23-1300

14 Garcia-Marin, R. et al. Prognostic and Therapeutic Implications of Immune Classification by CD8(+) Tumor-Infiltrating Lymphocytes and PD-L1 Expression in Sinonasal Squamous Cell Carcinoma. Int J Mol Sci 22 (2021). 10.3390/ijms22136926

15 Gu, J. T. et al. Characterization of the tumor immune microenvironment of sinonasal squamous-cell carcinoma. Int Forum Allergy Rhinol 12, 39–50 (2022). 10.1002/alr.22867

16 Deprez, M. et al. A Single-Cell Atlas of the Human Healthy Airways. Am J Respir Crit Care Med 202, 1636–1645 (2020). 10.1164/rccm.201911-2199OC

17 dos Santos Silva, E., Ulrich, M., Doring, G., Botzenhart, K. & Gott, P. Trefoil factor family domain peptides in the human respiratory tract. J Pathol 190, 133–142 (2000). 10.1002/(SICI)1096-9896(200002)190:2<133::AID-PATH518>3.0.CO;2-B

18 Odell, I. D. et al. Epiregulin is a dendritic cell-derived EGFR ligand that maintains skin and lung fibrosis. Sci Immunol 7, eabq6691 (2022). 10.1126/sciimmunol.abq6691

19 Taylor, P. R. et al. Macrophage receptors and immune recognition. Annu Rev Immunol 23, 901–944 (2005). 10.1146/annurev.immunol.23.021704.115816

20 Augsten, M. et al. CXCL14 is an autocrine growth factor for fibroblasts and acts as a multi-modal stimulator of prostate tumor growth. P Natl Acad Sci USA 106, 3414–3419 (2009). 10.1073/pnas.0813144106

21 Sjoberg, E., Augsten, M., Bergh, J., Jirstrom, K. & Ostman, A. Expression of the chemokine CXCL14 in the tumour stroma is an independent marker of survival in breast cancer. Br J Cancer 114, 1117–1124 (2016). 10.1038/bjc.2016.104

22 Cereijo, R. et al. CXCL14, a Brown Adipokine that Mediates Brown-Fat-to-Macrophage Communication in Thermogenic Adaptation. Cell Metab 28, 750–763 e756 (2018). 10.1016/j.cmet.2018.07.015

23 Lai, X. et al. CXCL14 Protects against Polymicrobial Sepsis by Enhancing Antibacterial Functions of Macrophages. Am J Respir Cell Mol Biol 67, 589–601 (2022). 10.1165/rcmb.2022-0249OC

24 Kumar, A. et al. CXCL14 Promotes a Robust Brain Tumor-Associated Immune Response in Glioma. Clin Cancer Res 28, 2898–2910 (2022). 10.1158/1078-0432.CCR-21-2830

25 Pacini, L., Cabal, V. N., Hermsen, M. A. & Huang, P. H. EGFR Exon 20 Insertion Mutations in Sinonasal Squamous Cell Carcinoma. Cancers (Basel) 14 (2022). 10.3390/cancers14020394

26 Black, S. et al. CODEX multiplexed tissue imaging with DNA-conjugated antibodies. Nat Protoc 16, 3802–3835 (2021). 10.1038/s41596-021-00556-8

27 Goltsev, Y. et al. Deep Profiling of Mouse Splenic Architecture with CODEX Multiplexed Imaging. Cell 174, 968–981 e915 (2018). 10.1016/j.cell.2018.07.010

28 Lu, J., Chatterjee, M., Schmid, H., Beck, S. & Gawaz, M. CXCL14 as an emerging immune and inflammatory modulator. J Inflamm-Lond 13, 1 (2016). 10.1186/s12950-015-0109-9

29 Wente, M. N. et al. CXCL14 expression and potential function in pancreatic cancer. Cancer Lett 259, 209–217 (2008). 10.1016/j.canlet.2007.10.021

30 Westrich, J. A. et al. CXCL14 suppresses human papillomavirus-associated head and neck cancer through antigen-specific CD8(+) T-cell responses by upregulating MHC-I expression. Oncogene 38, 7166–7180 (2019). 10.1038/s41388-019-0911-6

31 Parikh, A. et al. Malignant cell-specific CXCL14 promotes tumor lymphocyte infiltration in oral cavity squamous cell carcinoma. J Immunother Cancer 8 (2020). 10.1136/jitc-2020-001048

32 Tian, H. Y., Liang, Q., Shi, Z. & Zhao, H. Exosomal CXCL14 Contributes to M2 Macrophage Polarization through NF-kappaB Signaling in Prostate Cancer. Oxid Med Cell Longev 2022, 7616696 (2022). 10.1155/2022/7616696

33 Williams, K. A. et al. A systems genetics approach identifies CXCL14, ITGAX, and LPCAT2 as novel aggressive prostate cancer susceptibility genes. Plos Genet 10, e1004809 (2014). 10.1371/journal.pgen.1004809

34 Halse, H. et al. Multiplex immunohistochemistry accurately defines the immune context of metastatic melanoma. Sci Rep 8, 11158 (2018). 10.1038/s41598-018-28944-3

35 Parra, E. R. et al. Immune cellular patterns of distribution affect outcomes of patients with non-small cell lung cancer. Nat Commun 14, 2364 (2023). 10.1038/s41467-023-37905-y

36 Shurin, G. V. et al. Loss of new chemokine CXCL14 in tumor tissue is associated with low infiltration by dendritic cells (DC), while restoration of human CXCL14 expression in tumor cells causes attraction of DC both in vitro and in vivo. J Immunol 174, 5490–5498 (2005). 10.4049/jimmunol.174.9.5490

37 Starnes, T. et al. The chemokine CXCL14 (BRAK) stimulates activated NK cell migration: implications for the downregulation of CXCL14 in malignancy. Exp Hematol 34, 1101–1105 (2006). 10.1016/j.exphem.2006.05.015

38 Yu, G. et al. Prostate cancer-induced endothelial-cell-to-osteoblast transition drives immunosuppression in the bone-tumor microenvironment through Wnt pathway-induced M2 macrophage polarization. P Natl Acad Sci USA 121, e2402903121 (2024). 10.1073/pnas.2402903121

39 Witte, A. et al. The chemokine CXCL14 mediates platelet function and migration via direct interaction with CXCR4. Cardiovasc Res 117, 903–917 (2021). 10.1093/cvr/cvaa080

40 Kouzeli, A. et al. CXCL14 Preferentially Synergizes With Homeostatic Chemokine Receptor Systems. Front Immunol 11, 561404 (2020). 10.3389/fimmu.2020.561404

41 Chang, T. M. et al. CXCL14 promotes metastasis of non-small cell lung cancer through ACKR2-depended signaling pathway. Int J Biol Sci 19, 1455–1470 (2023). 10.7150/ijbs.79438

42 Sjoberg, E. et al. A Novel ACKR2-Dependent Role of Fibroblast-Derived CXCL14 in Epithelial-to-Mesenchymal Transition and Metastasis of Breast Cancer. Clin Cancer Res 25, 3702–3717 (2019). 10.1158/1078-0432.CCR-18-1294

43 Al Hamwi, G. et al. Proinflammatory chemokine CXCL14 activates MAS-related G protein-coupled receptor MRGPRX2 and its putative mouse ortholog MRGPRB2. Commun Biol 7, 52 (2024). 10.1038/s42003-023-05739-5

44 Schurch, C. M. et al. Coordinated Cellular Neighborhoods Orchestrate Antitumoral Immunity at the Colorectal Cancer Invasive Front. Cell 182, 1341–1359 e1319 (2020). 10.1016/j.cell.2020.07.005

45 Takano, Y. et al. Spatially resolved gene expression profiling of tumor microenvironment reveals key steps of lung adenocarcinoma development. Nat Commun 15, 10637 (2024). 10.1038/s41467-024-54671-7

46 Zhai, L. et al. IDO1 in cancer: a Gemini of immune checkpoints. Cell Mol Immunol 15, 447–457 (2018). 10.1038/cmi.2017.143

47 Ruysseveldt, E., Martens, K. & Steelant, B. Airway Basal Cells, Protectors of Epithelial Walls in Health and Respiratory Diseases. Front Allergy 2, 787128 (2021). 10.3389/falgy.2021.787128

48 Menzl, I. et al. Loss of primary cilia occurs early in breast cancer development. Cilia 3, 7 (2014). 10.1186/2046-2530-3-7

49 Collinson, R. & Tanos, B. Primary cilia and cancer: a tale of many faces. Oncogene 44, 1551–1566 (2025). 10.1038/s41388-025-03416-x

50 Sleeboom, J. J. F. et al. The extracellular matrix as hallmark of cancer and metastasis: From biomechanics to therapeutic targets. Sci Transl Med 16, eadg3840 (2024). 10.1126/scitranslmed.adg3840

51 Bull, J. A. et al. Combining multiple spatial statistics enhances the description of immune cell localisation within tumours. Sci Rep 10, 18624 (2020). 10.1038/s41598-020-75180-9

52 Hao, Y. et al. Integrated analysis of multimodal single-cell data. Cell 184, 3573–3587 e3529 (2021). 10.1016/j.cell.2021.04.048

53 Korsunsky, I. et al. Fast, sensitive and accurate integration of single-cell data with Harmony. Nat Methods 16, 1289–1296 (2019). 10.1038/s41592-019-0619-0

54 Korotkevich, G., Sukhov, V. & Sergushichev, A. Fast gene set enrichment analysis. bioRxiv (2016). 10.1101/060012

55 Liberzon, A. et al. Molecular signatures database (MSigDB) 3.0. Bioinformatics 27, 1739–1740 (2011). 10.1093/bioinformatics/btr260

56 Bergen, V., Lange, M., Peidli, S., Wolf, F. A. & Theis, F. J. Generalizing RNA velocity to transient cell states through dynamical modeling. Nat Biotechnol 38, 1408–1414 (2020). 10.1038/s41587-020-0591-3

57 Street, K. et al. Slingshot: cell lineage and pseudotime inference for single-cell transcriptomics. BMC Genomics 19, 477 (2018). 10.1186/s12864-018-4772-0

58 Cherry, C. et al. Computational reconstruction of the signalling networks surrounding implanted biomaterials from single-cell transcriptomics. Nat Biomed Eng 5, 1228–1238 (2021). 10.1038/s41551-021-00770-5

59 Andreatta, M. & Carmona, S. J. UCell: Robust and scalable single-cell gene signature scoring. Comput Struct Biotechnol J 19, 3796–3798 (2021). 10.1016/j.csbj.2021.06.043

60 Efremova, M., Vento-Tormo, M., Teichmann, S. A. & Vento-Tormo, R. CellPhoneDB: inferring cell-cell communication from combined expression of multi-subunit ligand-receptor complexes. Nat Protoc 15, 1484–1506 (2020). 10.1038/s41596-020-0292-x

61 La Manno, G. et al. RNA velocity of single cells. Nature 560, 494–498 (2018). 10.1038/s41586-018-0414-6

62 Browaeys, R., Saelens, W. & Saeys, Y. NicheNet: modeling intercellular communication by linking ligands to target genes. Nat Methods 17, 159–162 (2020). 10.1038/s41592-019-0667-5

63 Gu, Z., Eils, R. & Schlesner, M. Complex heatmaps reveal patterns and correlations in multidimensional genomic data. Bioinformatics 32, 2847–2849 (2016). 10.1093/bioinformatics/btw313

64 Gu, Z., Gu, L., Eils, R., Schlesner, M. & Brors, B. circlize Implements and enhances circular visualization in R. Bioinformatics 30, 2811–2812 (2014). 10.1093/bioinformatics/btu393

65 Dayao, M. T. et al. Deriving spatial features from in situ proteomics imaging to enhance cancer survival analysis. Bioinformatics 39, i140–i148 (2023). 10.1093/bioinformatics/btad245

66 Surwase, S. S. et al. Highly Multiplexed Immunofluorescence PhenoCycler Panel for Murine Formalin-Fixed Paraffin-Embedded Tissues Yields Insight Into Tumor Microenvironment Immunoengineering. Lab Invest 105, 102165 (2025). 10.1016/j.labinv.2024.102165

