## Supplemental Figures for "scRNA-seq and genomics analyses reveal key mechanisms of inverted papilloma-associated sinonasal squamous cell carcinoma malignant transformation"

S1

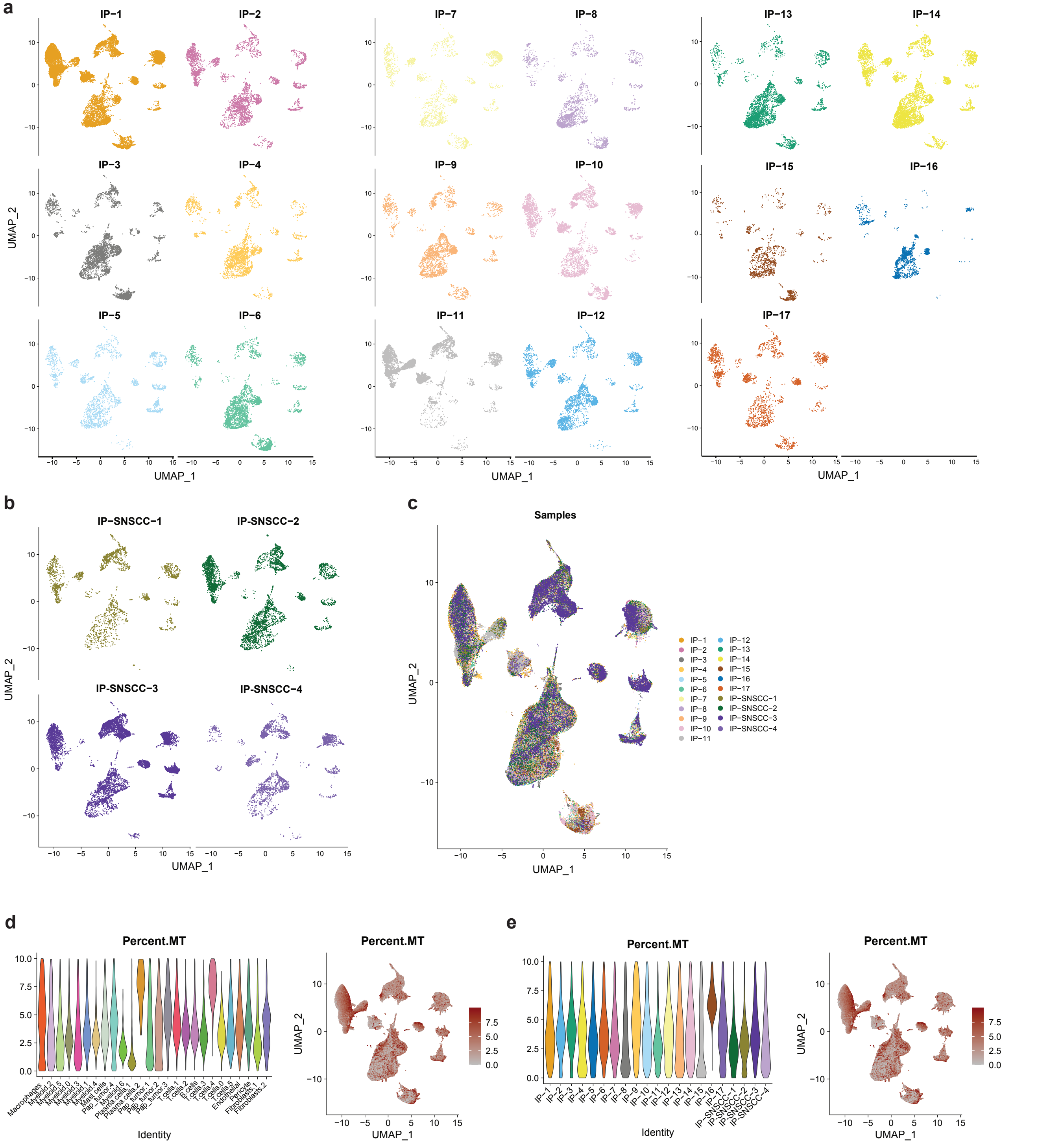


**Fig. S1**. Clusters are consistent across sampled tumors and do not represent patient-specific subpopulations. UMAP plot depicting cell clusters of individual IP (**a**), IP-SNSCC (**b**), and combined tumors (**c**) obtained from 19 patients. Violin plots with corresponding UMAP plots demonstrating the percentage of reads that map to the mitochondrial genome for each cluster (**d**) and for each individual patient (**e**).

S2

**a**

| Markers | Scores Sinonasal IP | Scores SCC |
|---|---|---|
| CD8 | 2 | 2 |
| CD20 | 2 | 2 |
| CD4 | 2 | 2 |
| HLA-A | 2 | 1 |
| CD14 | 2 | 2 |
| CD68 | 2 | 2 |
| Ki67 | 3 | 3 |
| SMA | 2 | 2 |
| HLA-DR | 2 | 2 |
| CD44 | 2 | 1 |
| Pan-Cytokeratin | 2 | 2 |
| CD3e | 2 | 2 |
| CD45RO | 1 | 1 |
| CD45 | 2 | 2 |
| Granzyme B | 0 | 0 |
| FOXP3 | 1 | 2 |
| E-cadherin | 2 | 2 |
| Collagen IV | 2 | 3 |
| Podoplanin | 2 | 1 |
| PD-1 | 0 | 1 |
| CD38 | 2 | 2 |
| PD-L1 | 2* | 2 |
| IFNG | 2 | 2 |
| IDO1 | 2 | 2 |
| CD107a | 2 | 2 |

| Score | Interpretation | Include in analysis |
|---|---|---|
| -1 | non-specific binding | no |
| 0 | no signal | no |
| 1 | weak signal | no |
| 2 | good staining | yes |
| 3 | saturated | yes |

**b**

**c**

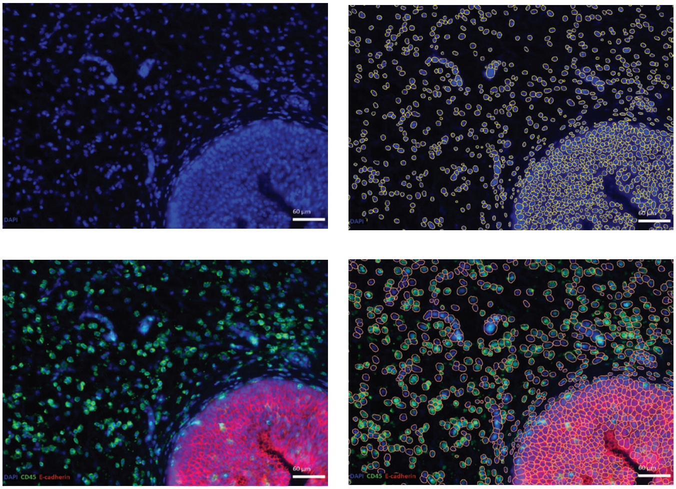

**d**

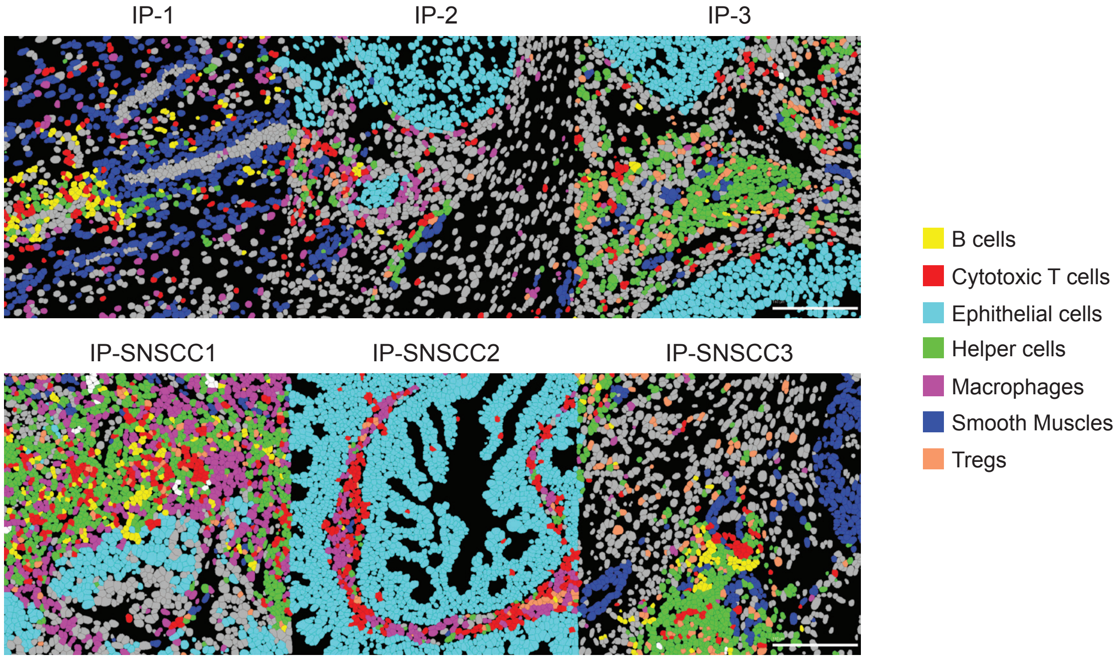


**Fig. S2**. **Performance of single cell segmentation algorithms.** (**a**) Quality control performance score for each marker utilized. StarDist single cell segmentation delineating (**b**) nuclear marker, DAPI and (**c**) cell membrane markers CD45 and E-cadherin in sample IP tissue. (**d**) sample supervised phenotyping results using lineage marker expression in individual IP (n = 3) and IP-SNSCC (n = 3) tissue samples. Scale bar = 60µm

S3

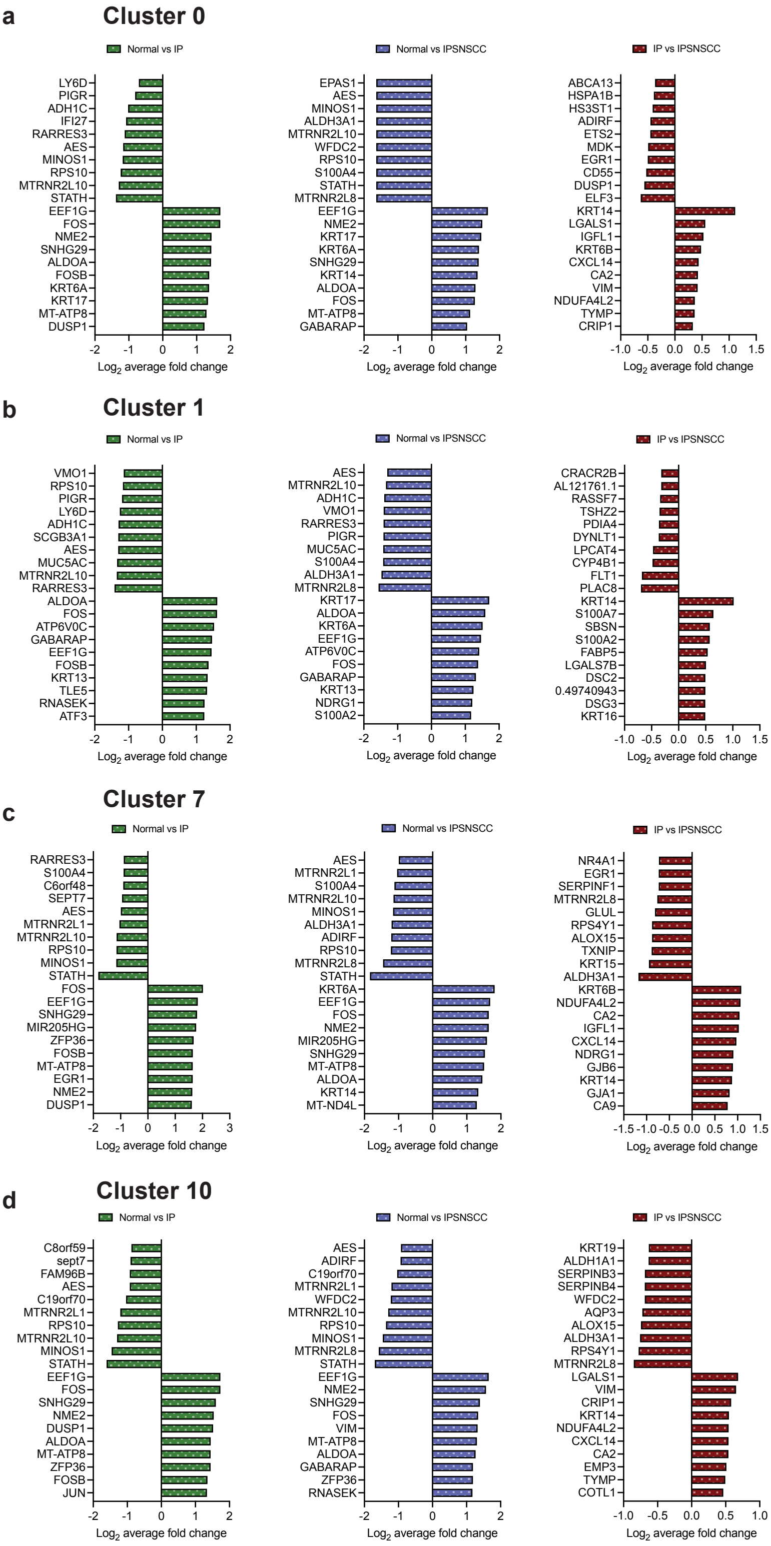


**Fig. S3. Defining features of identified clusters in pseudotime trajectory analysis.** Differential gene expression (top ten upregulated and downregulated) between healthy control (normal) and IP, normal and IPSNSCC, IP and IPSNSCC in (**a**) Cluster 0, (**b**) Cluster 1, (**c**) Cluster 7, and (**d**) Cluster 10.

S4

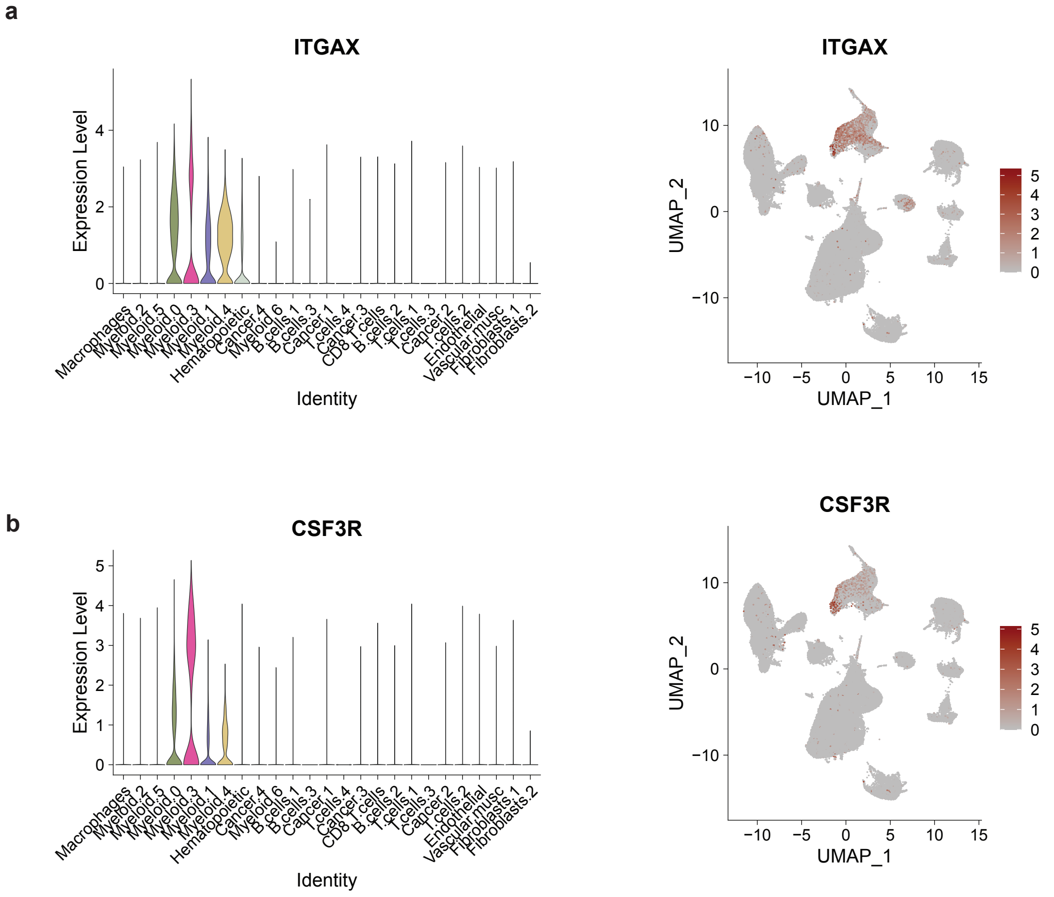


**Fig. S4. Dendritic and granulocytic markers are in a distinct cell cluster.** Expression levels and UMAP plots of (**a**) *ITGAX* and (**b**) *CSF3R* by cell cluster.

S5

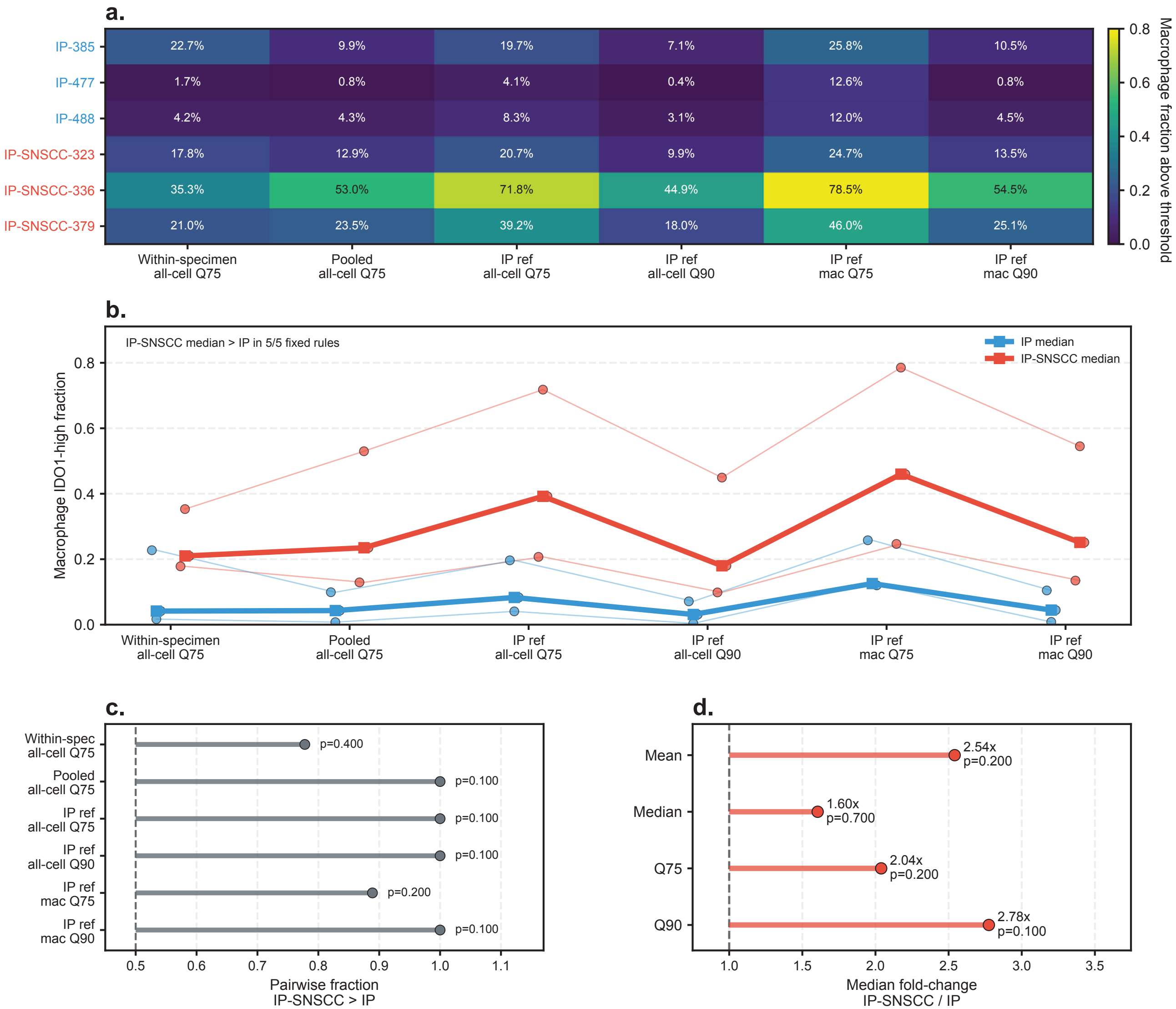


**Fig. S5. Sensitivity of macrophage IDO1-high fraction to threshold definition.** Macrophage IDO1-high fraction was recalculated across alternative intensity thresholds to assess whether the IP-SNSCC versus IP cohort pattern depended on the within-specimen all-cell Q75 threshold used in the primary analysis. **a.** Heatmap of the percentage of macrophages classified as IDO1-high in each specimen under each threshold rule. The within-specimen all-cell Q75 rule defines the cutoff as the 75th percentile of IDO1 intensity among all cells in the same specimen, then applies that cutoff to macrophages from that specimen. Fixed sensitivity thresholds use pooled all-cell Q75, IP-reference all-cell Q75/Q90, and IP-reference macrophage Q75/Q90. **b.** Sample-level macrophage IDO1-high fractions across threshold definitions. Thin lines trace individual specimens, and bold lines show cohort medians. The IP-SNSCC median exceeded the IP median across all five fixed-threshold sensitivity rules. **c.** Directional robustness summarized as the fraction of all IP-SNSCC-versus-IP sample pairs in which the IP-SNSCC value was higher for each threshold rule. The dashed reference line marks 0.5, corresponding to no directional preference. P values are exact two-sided Mann-Whitney U tests. **d.** Threshold-free continuous macrophage IDO1 summaries plotted as median fold-change for IP-SNSCC versus IP. Metrics include macrophage mean IDO1, median IDO1, Q75, and Q90. P values are exact two-sided Mann-Whitney U tests.

S6

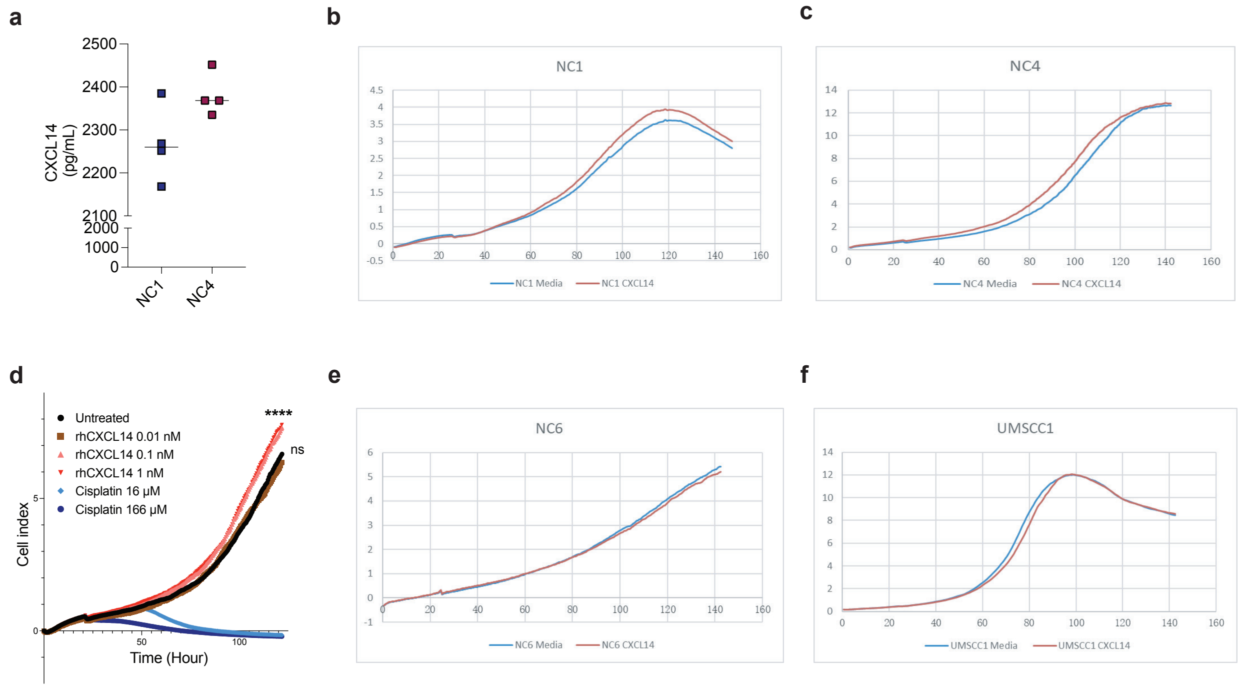


**Fig. S6. CXCL14 is not a direct modulator of IP-SNSCC proliferation.** (**a**) CXCL14 expression in untreated NC1 and NC4 cell lysates. Cell proliferation as determined by cellular impedance for (**b**) NC1 sinonasal squamous cell carcinoma (SNSCC)-derived cell line, and (**c**) NC4 IPSNSCC-derived cell line treated with control (vehicle) or rhCXCL14 50 ng/mL (**d**) Dose-dependent treatment in NC4 IPSNSCC-derived cell line. Cell proliferation as determined by cellular impedance for (**e**) NC6 SNSCC-derived cell line, and (**f**) UMSCC1 non-sinonasal squamous cell carcinoma-derived cell line treated with control (vehicle) or rhCXCL14 50 ng/mL. ****$p < 0.0001$; ns = not significant ($p > 0.05$). Statistical comparisons performed by two-way ANOVA with Tukey's multiple comparisons test. Each plot is indicative of at least 2 independent experiments.

S7

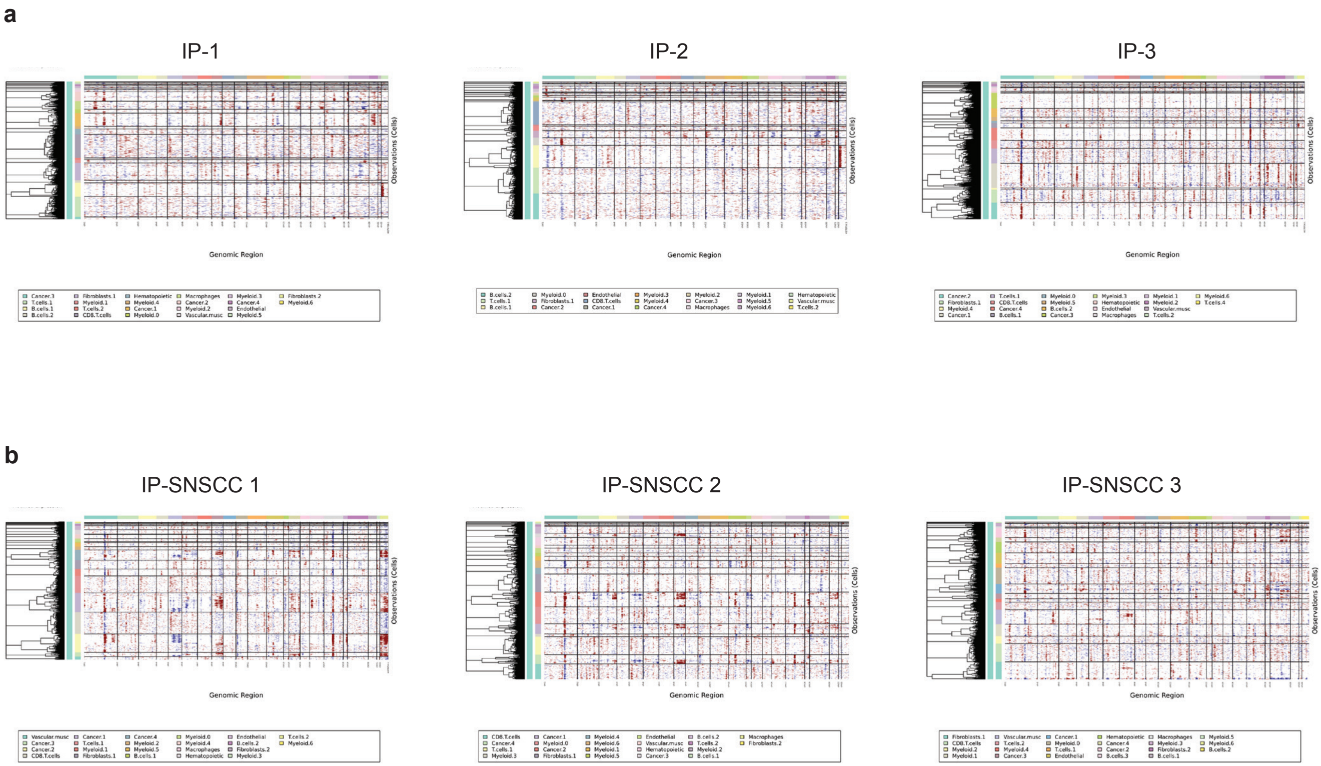


**Fig. S7. Characterization of copy number alterations inferred from scRNA-seq data.** InferCNV heatmap depicting large-scale chromosomal copy-number variations (CNVs) at single cell resolution in representative (**a**) IP tumors and (**b**) IPSNSCC tumors. Rows: Cell cluster; Columns: Chromosome position. Red: Amplifications; Blue: Deletions.

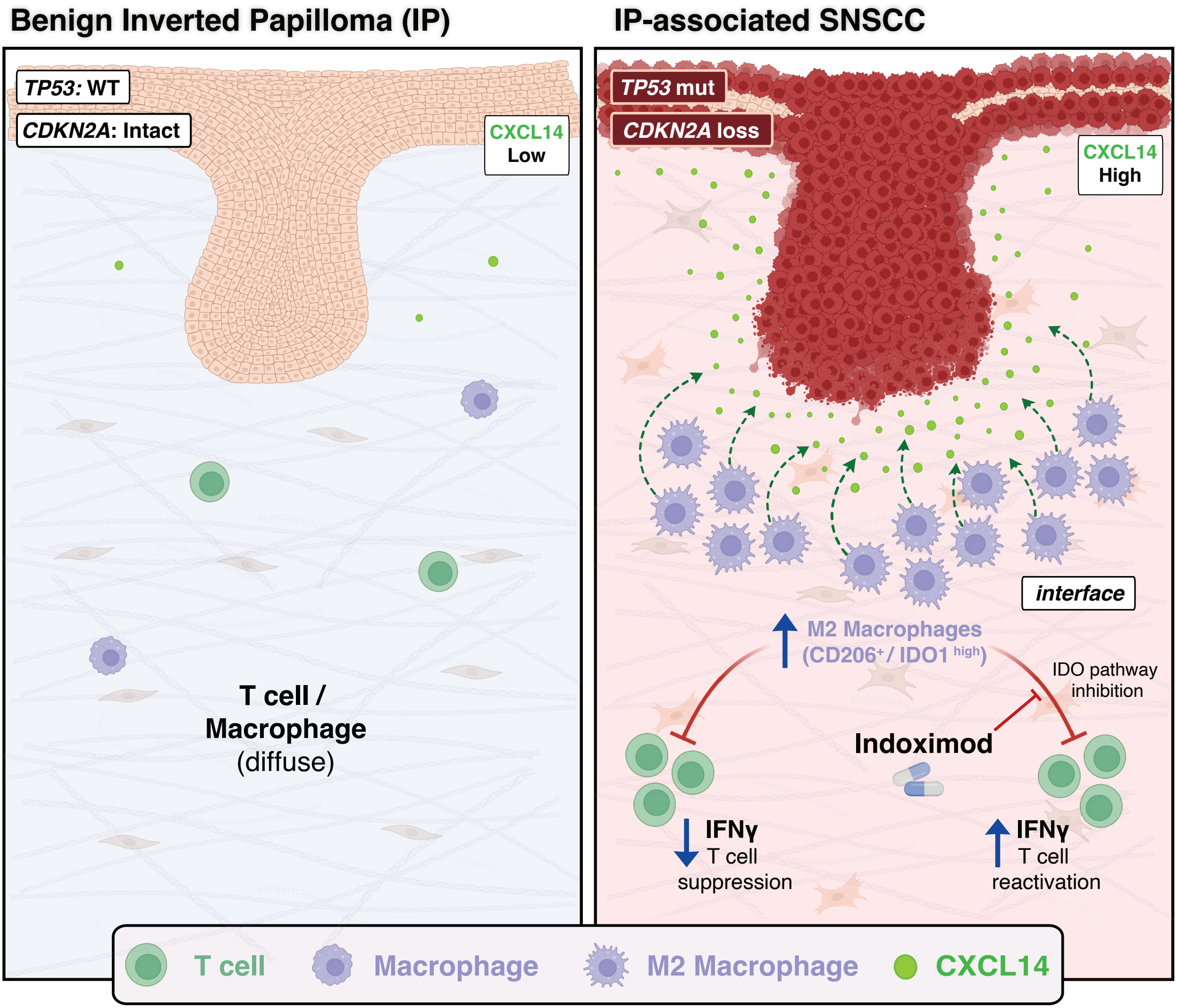


**Fig. S8. Schematic model of the CXCL14-IDO immunosuppressive program associated with malignant transformation from IP to IP-associated SNSCC.** In benign inverted papilloma (**IP, left**), the lesional epithelium retains wild-type *TP53* and intact *CDKN2A* status and shows low *CXCL14* expression. The immune microenvironment is characterized by diffusely distributed T cells and macrophages that remain spatially distant from the lesional epithelium. In IP-associated sinonasal squamous cell carcinoma (**IP-SNSCC, right**), malignant progression is associated with recurrent *CDKN2A* loss and *TP53* mutations, accompanied by increased tumor-cell *CXCL14* expression and secretion. Tumor-derived CXCL14 promotes recruitment and immunoregulatory polarization of monocytes/macrophages at the tumor-stroma interface, leading to enrichment of $CD206^{+}$/IDO1-high macrophages. These interface-localized IDO1-expressing macrophages suppress T-cell effector function, reflected by reduced IFNγ production. Pharmacologic inhibition of the IDO pathway with Indoximod attenuates this suppressive program and restores T-cell IFNγ production.
